# The stem cell compartment in human oral mucosa and its activation in oral lichen planus

**DOI:** 10.64898/2026.03.04.709521

**Authors:** Olaf J. F. Schreurs, Stefano Fedele, Stephen Porter, Gry K. Kjølle, Karl Schenck, Tine M. Søland, Gernot Walko

**Affiliations:** Institute of Oral Biology, Dental Faculty, University of Oslo, Oslo, Norway; UCL Eastman Dental Institute, University College London, London, UK; NIHR UCLH Biomedical Research Centre, London, UK; Oris Dental Hamar, Hamar, Norway; Department of Pathology, Oslo University Hospital, Oslo, Norway; Centre for Oral Immunobiology and Regenerative Medicine, Institute of Dentistry, Queen Mary University of London, London, UK

**Keywords:** Tissue-resident stem cells, oral mucosa, oral lichen planus, keratin intermediate filaments

## Abstract

In mice, oral epithelial stem cells (OESCs) are essential for oral mucosal homeostasis and repair. Less is known regarding the role of OESCs in the human oral mucosa. Here, we studied the behaviour of OESCs and their contribution to tissue maintenance and repair in oral lichen planus (OLP). OLP is a chronic T cell-mediated disease characterized by basal keratinocyte degeneration, epithelial atrophy, acanthosis, and hyperkeratosis. Using immunohistological techniques and semi-automated image analysis, we observed that in OLP proliferative activity was increased in the normally largely quiescent basal cell compartment. In areas of OLP mucosa with intact basal cell layer, expression of NGFR, KRT15, and KRT19–markers of slowly cycling ‘reserve’ OESCs, was strongly reduced or absent. In contrast, expression of CSPG4, a marker for actively cycling stem cells, was increased in OLP basal cells. Tissue compartmentalization, as evaluated by keratin expression, was strongly disturbed. Taken together, our findings indicate that the inflammation in OLP leads to activation and proliferation of OESCs that give rise to a population of cells with an aberrant differentiation programme. Along with the well-documented epithelial up-regulation of anti-apoptotic proteins in OLP, this likely reflects an attempt by the epithelium to avoid overt ulceration.

## Introduction

The stratified squamous epithelia of the skin and the body’s mucosal surfaces protect the underlying tissue from exposure to harmful agents, pathogenic microorganisms and physical forces (Jones and Klein, 2013, Maurizi et al., 2021, Pereira and Sequeira, 2021). These epithelial tissues are formed of tightly packed epithelial cells (keratinocytes) including a basal cell layer that rests on the subepithelial connective tissue. Stratified squamous epithelia display a high tissue turnover and the capacity of self-renewal. The most terminally differentiated epithelial cells are continually sloughed off at the epithelial surface, and to ensure tissue homeostasis, this cell loss is replaced by new cells that are generated in the lower parts of the epithelium. While the terminally differentiating cells move towards the epithelial surface, they undergo different stages of programmed morphological and biochemical changes, tailored to fulfill the protective functions required by the squamous epithelium (Fuchs, 2008, Pereira and Sequeira, 2021, Zhang et al., 2021).

Research in mice has shown that the maintenance of homeostasis and tissue repair in stratified squamous epithelia is driven by tissue-resident stem cells, and multiple models have been proposed to explain the mechanisms by which stem cells sustain tissue homeostasis (Cockburn et al., 2022, Pereira and Sequeira, 2021). In the oral cavity of mice, distinct populations of oral epithelial stem cells (OESCs) appear to maintain the different types of oral epithelium (Byrd et al., 2019, Jones et al., 2019). The OESCs may also be involved in mucosal carcinogenesis (Jones and Klein, 2013).

The dogmatic view of stratified squamous epithelial homeostasis is that proliferation is confined to the basal cell layer. In this scenario, as is seen in murine stratified squamous epithelia, quiescent and actively cycling stem cells coexist with transit-amplifying progenitor cells (TACs), a cell population with a high but limited proliferative capacity that is committed to terminal differentiation, within this layer. This cellular intermingling may complicate the characterization of squamous epithelial stem cells in the mouse. In contrast, in human stratified squamous epithelia such as the oral mucosa, ectocervix, and vagina, most cells expressing Ki-67 and other proliferation markers are found in the parabasal cell layers, while basal cell proliferation is less frequent (Andl et al., 2016, Dunaway et al., 2019, Harris et al., 2025, Karatsaidis et al., 2003b). Recent studies on human epidermis have also noted a similar compartmentalization of proliferative activity, at least in certain skin areas (Noske et al., 2016, Pontiggia et al., 2022, Wang et al., 2020, Wiedemann et al., 2023). Together, this body of evidence has led to the view that the basal cell compartment in stratified squamous epithelia of long-lived species (including human) hosts a population of largely quiescent and only slowly cycling ‘reserve’ stem cells that only become significantly activated in response to devastating events that kill a large proportion of epithelial cells, such as exposure to irradiation, or in high risk potential malignant disorders (Dunaway et al., 2019). In contrast, the proliferative parabasal compartment acts as the main driver of tissue maintenance.

Oral lichen planus (OLP) is a commonly occurring T cell-mediated chronic inflammatory disease of the oral mucosa displaying profound changes in the epithelium of the affected areas, with an overall estimated pooled prevalence of 0.89 % (Li et al., 2020). Clinically, OLP occurs in quiescent and active forms that can vary with time (Andreasen, 1968, Silverman et al., 1985, Thorn et al., 1988), and the morphological appearance is described as reticular (RET-OLP), papular, plaque-like, atrophic (erythematous), erosive (ulcerative), and rarely as bullous (Cheng et al., 2016, Ingafou et al., 2006). Atrophic (ATR-OLP; also termed erosive) and ulcerative OLP constitute the more tissue-destructive disease forms, while purely reticular (RET-OLP), papular, and plaque-like forms are associated with few symptoms, no atrophy or loss of the epithelium, and therefore are considered to represent a less active disease phase (Carrozzo et al., 2019). Although the buccal mucosa is the most frequently affected oral site, all mucosal areas of the oral cavity can be involved, including the gingiva. Concomitant involvement of extraoral sites (cutaneous, genital, scalp, nail, oesophageal, or ocular lichen planus) is not notably common (Bidarra et al., 2008, Eisen, 1999, Feldmeyer et al., 2020, Schlosser, 2010). Oral lichen planus etiopathogenesis remains unclear, however in some patients triggers can be identified and include local reactions to dental restorative materials (OLCL, oral lichenoid contact lesion) or medication (oral lichenoid drug reaction (OLDR), graft versus host disease (GvHD) (Carrozzo et al., 2019), or the loss of immune tolerance associated with immune checkpoint inhibitors (Sehbai et al., 2022). Histologically, the disease is characterized by basal epithelial cell damage, epithelial atrophy, and a dense inflammatory infiltrate in the connective tissue of the lamina propria, primarily consisting of T-lymphocytes (Andreasen, 1968, Hedberg et al., 1986, Odell, 2025). Both antigen-specific and non-specific pathogenic mechanisms have been proposed to cause the local destruction and epithelial remodelling in OLP (Conrad and Gilliet, 2012, Maciel et al., 2025, Roopashree et al., 2010, Thornhill, 2001). The former mechanism includes antigen presentation by basal keratinocytes, activation of CD4+ helper T cells, cytokine release, and keratinocyte cell death induced by cytotoxic CD8+ T cells. Non-specific mechanisms can act through activation of Toll-like receptors on basal keratinocytes (Janardhanam et al., 2012) or release of pro-inflammatory mediators and proteases by e.g. mast cells in OLP lesions, resulting in T-cell infiltration, disruption of the basement membrane and intrusion of immune cells into the epithelium (Roopashree et al., 2010). While most focus has been on apoptosis as the form of cell death that is induced in the basal epithelial cells in OLP (Maciel et al., 2025), compelling recent information indicates that other cell death programs such as pyroptosis (Yang et al., 2024) and necroptosis (Kurzen et al., 2025) also can occur.

Independently of the form of OLP, the epithelium is always affected (Schreurs et al., 2020a, Schreurs et al., 2020b), and total epithelial loss, i. e. ulceration, is relatively seldom (Ingafou et al., 2006). How the surviving epithelial cells with proliferative capacity deal with the tissue damage and contribute to epithelial maintenance and repair in OLP remains insufficiently investigated. In this study, we therefore have set out to investigate the hypothesis that the otherwise quiescent OESCs in the basal cell compartment of OLP become activated by the altered signalling environment. To this end, we comprehensively profiled the expression of markers specific for the different epithelial tissue compartments, as well as markers of proliferation and OESC activation, in the oral mucosa of an OLP patient cohort. We found that there is considerable epithelial tissue remodelling in OLP, which is linked to OESC activation, increased cell proliferation, and aberrant terminal differentiation.

## Results

### Increased proliferation in the basal cell compartment of oral mucosa in OLP

To test our hypothesis that the tissue damage in OLP triggers activation of the OESCs within the basal compartment of the oral mucosa, we first compared the location of proliferating cells in oral mucosa biopsies taken from individuals with healthy tissue (normal oral mucosa, NOM) and from patients with reticular OLP (RET-OLP) and atrophic OLP (ATR-OLP) by examining the expression of the proliferation marker Ki-67 (Figure 1). Ki-67 protein is present during all active phases of the cell cycle (G_1_, S, G_2_, and mitosis) and is absent in resting (quiescent) cells (G_0_). In agreement with previous observations (Andl et al., 2016), visual inspection showed that the majority of Ki-67-positive cells in NOM was found in the first parabasal cell layer (Figure 1A). The numbers of proliferating cells were increased in the basal cell compartment of histologically intact epithelium in RET-OLP (Figure 1B) and ATR-OLP (Figure 1C) oral mucosa. Of note, in ATR-OLP tissue, Ki-67-positive cells were also detectable in surviving basal cells in areas of basal cell layer degeneration (Figure 1D). Expression of p21 (CDKN1A), which in squamous epithelia is involved in cell cycle withdrawal and terminal differentiation commitment (de Pedro et al., 2021, Topley et al., 1999), was largely confined to cells above the basal cell layer in both NOM and OLP (Supplementary Figure S1). This agrees with the concept that the parabasal cells in the oral mucosa constitute a population of TACs. In OLP, basal p21^+^ cells appeared more frequently and could be arrested in cell-cycle (Bascones-Ilundain et al., 2007)(Supplementary Figure S1Bii, Cii). Semi-automated quantification of Ki-67-stained cells across large areas of epithelial tissue displaying recognisable basal membrane, confirmed that there were significantly more proliferating basal cells and an increase of proliferative activity in the basal cell compartment in RET-OLP and ATR-OLP as compared with NOM (Figure 1E). These observations provide evidence for increased activation and proliferation of ‘reserve’ OESCs in the basal cell compartment of OLP oral mucosa.

**Figure 1.**
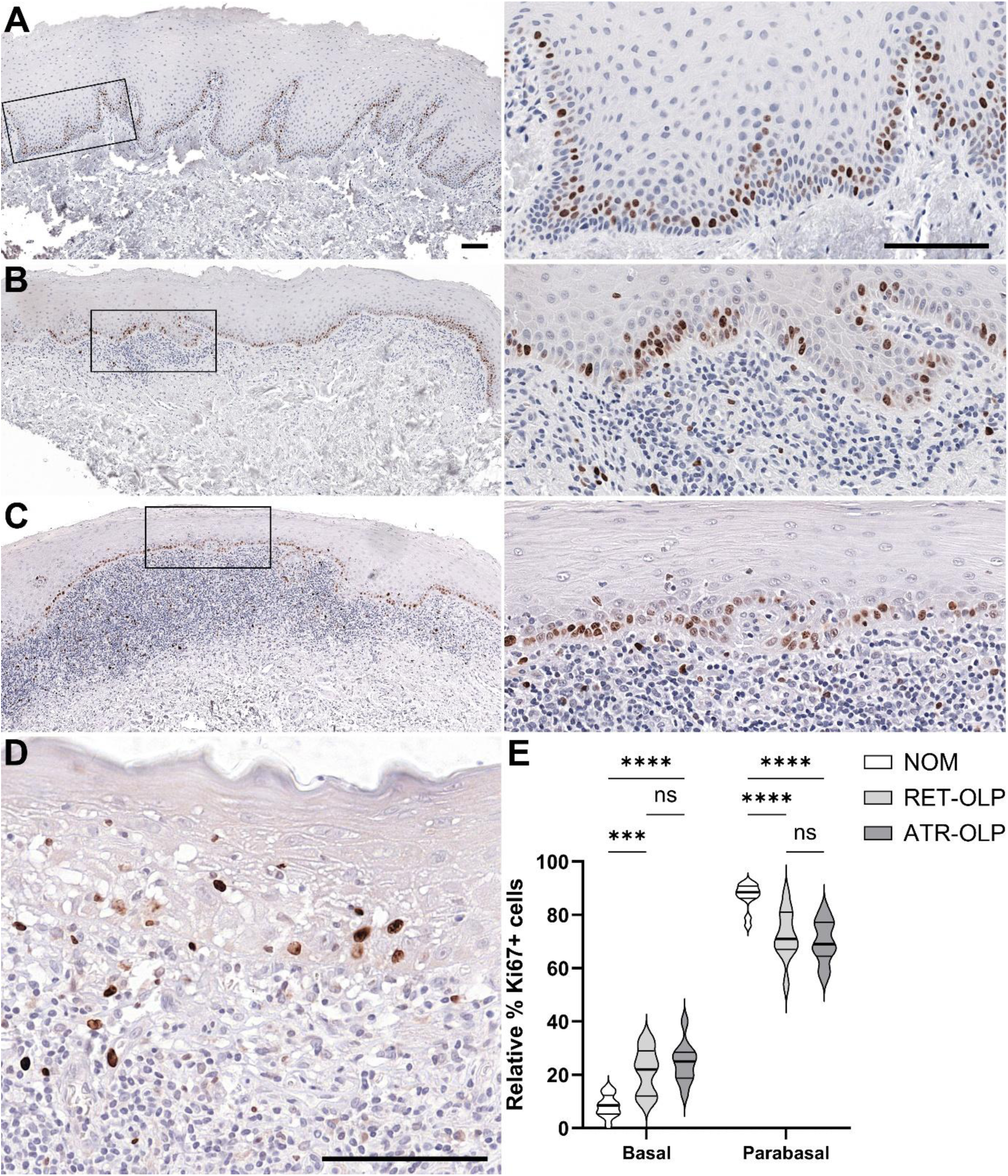
Increased proliferation of basal cells in OLP. Expression of proliferation marker Ki-67 in normal oral mucosa (NOM; A), reticular OLP (RET-OLP; B), and atrophic OLP (ATR-OLP; C, D). Bars, 100 µm. (E) Quantification of Ki-67-positive epithelial cells in basal versus parabasal tissue layers (number of tissue samples: n=12, NOM; n=11, RET-OLP; n=10, ATR-OLP). Violin plots show median (thick line) and upper- and lower quartiles (thin lines). p-values are shown for multiple comparisons, two-way ANOVA and Šidak’s multiple comparison test. ns, p=0.1234, *p=0.0332; **p=0.0021; ***p=0.0002; ****p<0.0001.

### Oral mucosal epithelial tissue compartmentalisation is markedly disturbed in OLP

In NOM, the different tissue compartments that accommodate quiescent, proliferating, and terminally differentiating cells are respectively characterized by a distinctive expression pattern of keratin intermediate filament proteins. Keratins not only provide mechanical cell stability and integrity, but also participate in various regulatory functions, including cell signalling, cell cycle regulation, and cell death (Pan et al., 2013, Redmond et al., 2026). The basal cells in NOM expressed the OESC markers keratins (K)15 (Andl et al., 2016) (Figure 2A) and K19 (Schreurs et al., 2020b) (Figure 2D), as well as K8 (Figure 2G) (Schreurs et al., 2020b), K5 (Figure 2J), and K14 (Figure 2M). Very few parabasal cells were positive for K19 (Figure 2D), while many cells in the first parabasal cell layer expressed K15, K8, and K14 (Figure 2A, G, M). In NOM, K5 was most strongly expressed in parabasal cells, while suprabasal cells were weakly positive for K5 (Figure 2J). We found that only very few individual cells stained positive for K15 and K8 in ATR-OLP oral mucosa (Figure 2C, I), while we observed larger patches of basal cells still expressing these keratins in RET-OLP oral mucosa (Figure 2B, H). K19 expression was completely lost in both RET-OLP and ATR-OLP (Figure 2E, F). In contrast to NOM, K5/14 expression expanded to the suprabasal cell layers in both RET-OLP and ATR-OLP (Figure 2K, L, N, O). Expression of the stress- and wound-activated K16, which is already uniquely present in NOM in contrast to healthy epidermis (Iglesias-Bartolome et al., 2018, Schreurs et al., 2020b), was predominantly restricted to non-proliferating cells above the parabasal cell layer (Figure 2P). Interestingly, in RET-OLP and ATR-OLP buccal mucosa, K16 expression was already apparent in most parabasal cells and some basal cells (Figure 2Q, R). As is typical for a stratified non-keratinizing epithelium, in NOM all suprabasal terminally differentiating cells (excluding the parabasal cells) expressed K4 (Boisnic et al., 1995) (Figure 2S). In agreement with other studies (Boisnic et al., 1995, Danielsson et al., 2014), we found expression of K4 to be strongly reduced or absent in the suprabasal cell layers of OLP oral mucosa (Figure 2T, U). By analysing published microarray data (Danielsson et al., 2014), we found that the observed changes in cytokeratin protein expression between NOM and OLP epithelial tissue were detectable also at the mRNA level, featuring downregulation of *KRT4*, -*8*, -*15*, and -*19*, and upregulation of *KRT6A*, -*6B*, and -*16* in OLP mucosa (Supplementary Figure S2A, B). Intriguingly, *KRT1* and *KRT10*, differentiation markers of keratinized squamous epithelia (Pereira and Sequeira, 2021), were upregulated in OLP mucosa, as were genes belonging to the epidermal differentiation complex (EDC) (Kypriotou et al., 2012) (Supplementary Figure S2B, C). In NOM buccal mucosa, K10 was either absent or weakly expressed in suprabasal cells (Figure 2Vi, Vii). In contrast, we noticed moderately to strongly K10-expressing suprabasal cells in RET-OLP and ATR-OLP (Figure 2Wi, Wii, Xi). In some cases of ATR-OLP, K10 was expressed throughout the epithelium (Figure 2Xii). The observed changes in keratin expression pattern are summarized in Supplementary Table S1.

**Figure 2.**
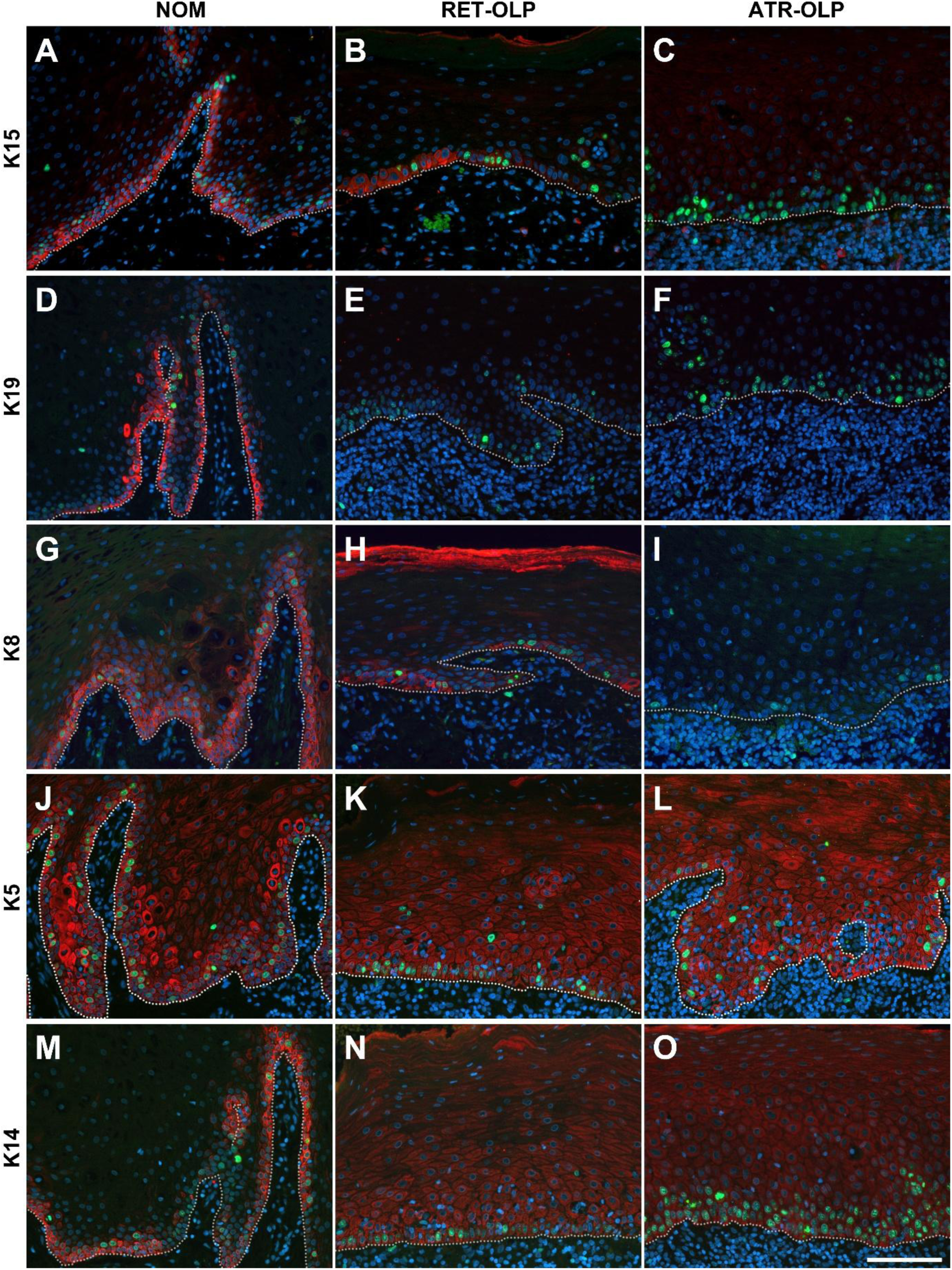
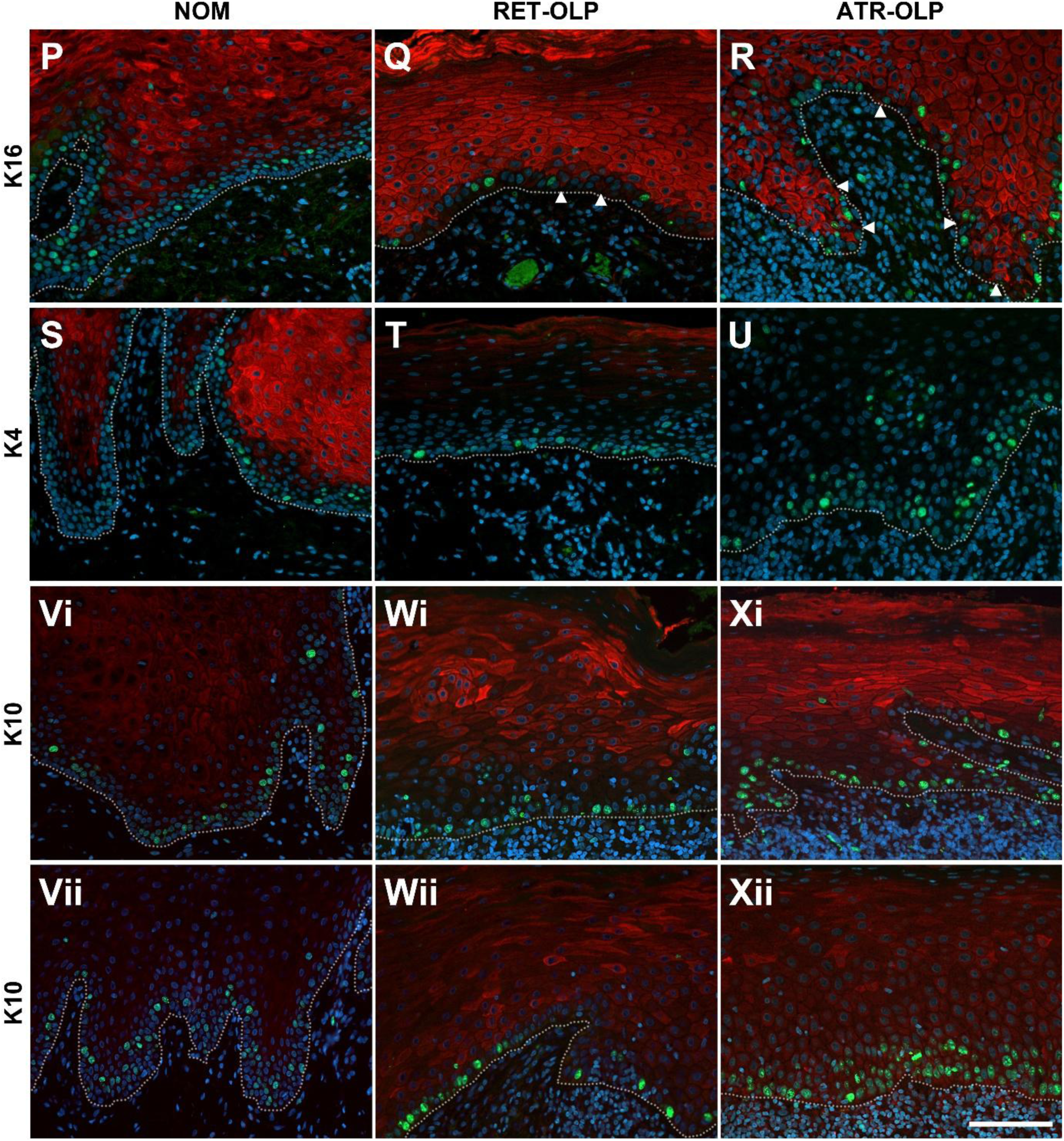
Altered mucosal tissue compartmentalization in OLP. Expression of major keratin intermediate filament proteins (red colour) in normal oral mucosa (NOM; A, D, G, J, M, P, S, Vi, Vii), reticular OLP (RET-OLP; B, E, H, K, N, Q, T, Wi, Wii), and atrophic OLP (ATR-OLP; C, F, I, L, O, R, U, Xi, Xii). Representative images show expression of keratin (K)15 (A–C), K19 (D–F), K8 (G–I), K5 (J–L), K14 (M–O), K16 (P–R), K4 (S–U), and K10 (Vi–Xii). Tissue sections were also labelled for Ki-67 (green) and counterstained with DAPI (blue). Dashed lines indicated the boundary between epithelium and underlying stromal tissue. Bars, 50 µm.

Together with our observation of increased basal cell proliferation, these findings highlight drastic alterations of tissue compartmentalisation and homeostasis in OLP mucosal epithelium that include loss of the parabasal (Ki-67^+^/K4^-^/K16^-^/K19^-^) cell compartment, increase of proliferative activity to the basal cell compartment, and aberrant terminal differentiation towards a keratinized squamous epithelium.

### Expression of marker proteins associated with OESCs are altered in OLP

It was previously suggested that OESCs in the basal cell compartment of NOM are kept in a quiescent state through TGFβ-SMAD signalling (Andl et al., 2016). This signalling axis was reported to be partially disrupted in OLP (Karatsaidis et al., 2003a) and the presently observed shift of proliferative activity to the basal cell compartment in OLP indicates increased activation of the otherwise slowly cycling ‘reserve’ OESCs. To provide evidence in support of this hypothesis, we analysed the expression of proteins related to OESCs in oral mucosa of NOM versus OLP.

Consistent with the concept that immune cell attack on the basal cell compartment in OLP does not lead to complete loss of all OESCs, we found that expression of the stemness-associated protein integrin (ITG)B1, a key components of focal adhesions (Gunnarsson et al., 2017, Jones and Watt, 1993), was preserved in RET-OLP and ATR-OLP tissue areas with no apparent epithelial damage (Figure 3A–C). Compared to NOM, in RET-OLP the basal cell surface staining of the hemidesmosome component Collagen XVII (COL17A1, previously called BP180) was more patchy (Figure 3D, E), while in ATR-OLP it was mostly absent from the basal cell surface and basal cells instead displayed circumferential membranous and/or cytoplasmic staining (Figure 3F), indicating deficient hemidesmosome homeostasis as noted previously (Schreurs et al., 2020a). However, such mild deficiencies in hemidesmosome assembly are not enough to cause epithelial detachment as also observed in studies focussing on the epidermis (Ackerl et al., 2007, Liu et al., 2019).

**Figure 3.**
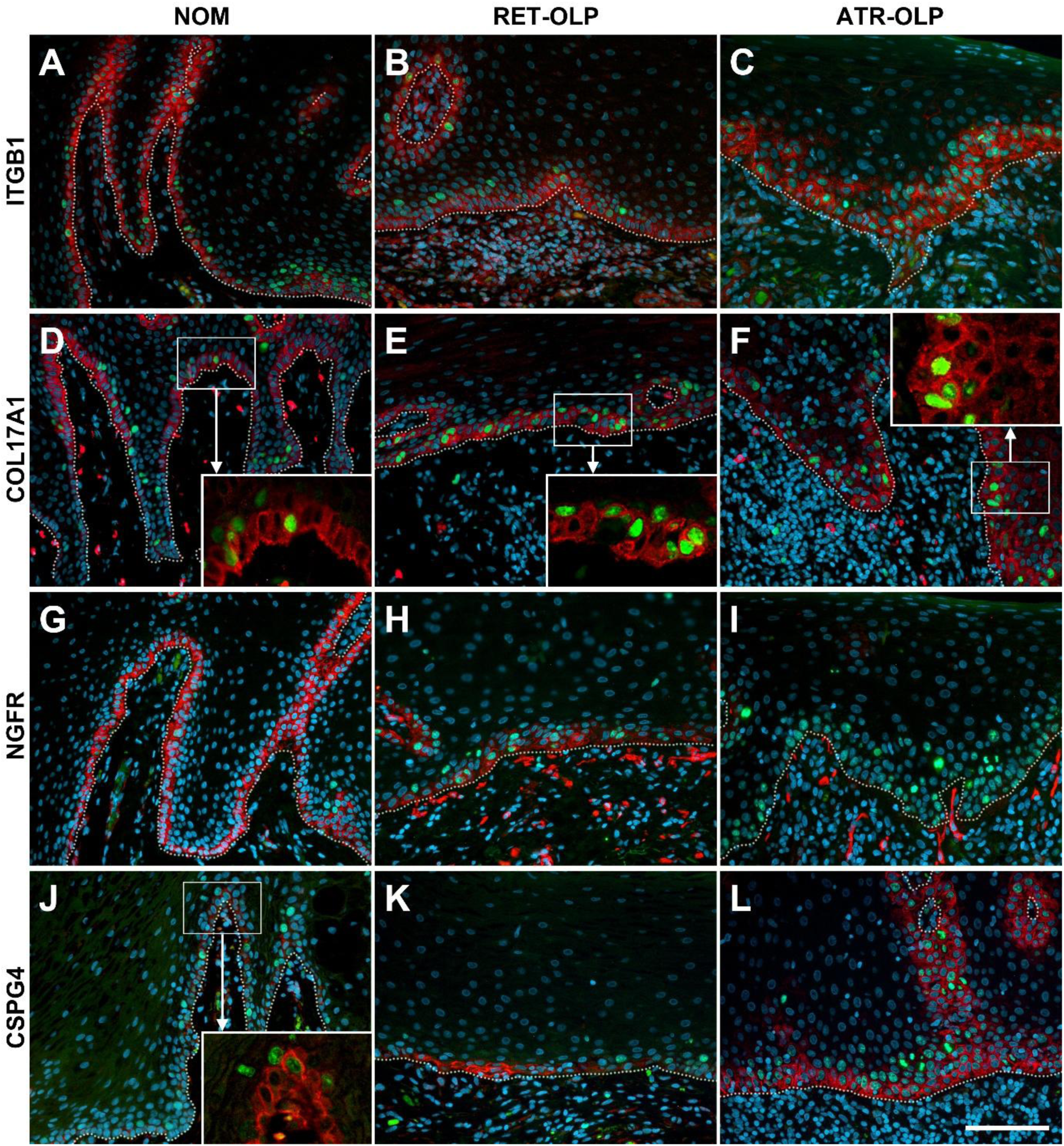
Altered expression of OESC markers in OLP. Expression of protein markers (red colour) of OESCs in normal oral mucosa (NOM; A, D, G, J), reticular OLP (RET-OLP; B, E, H, K), and atrophic OLP (ATR-OLP; C, F, I, L). Representative images show expression of ITGB1 (A–C), Collagen XVII (COL17A1, D–F), NGFR (G–I), CSPG4 (J–L). Tissue sections were also labelled for Ki-67 (green) and DAPI (blue). Dashed lines indicated the boundary between epithelium and underlying stromal tissue. Bar, 50 µm.

The low affinity neurotrophin receptor p75 (p75NTR, NGFR) labels quiescent cells in the basal cell compartment of NOM which, upon isolation from the tissue, become highly proliferative in culture (Nakamura et al., 2007) – a cellular behaviour that is also typical for stem cells sorted from human epidermis on the basis of high ITGA6 and low CD71 surface expression (Li and Kaur, 2005, Schluter et al., 2011). In NOM, NGFR was exclusively expressed in the basal cell compartment, with a tendency towards strongest expression in basal cells at the top of the papillary lamina propria (i. e. closest to the mucosal tissue surface) (Figure 3G, Table 1). In contrast, in RET-OLP and ATR-OLP fewer basal cells expressed NGFR and in some ATR-OLP tissue sections NGFR staining was largely undetectable (Figure 3H, I; Table 1). Of note, the loss of NGFR expression in OLP paralleled that of KRT15 (Figure 2A–C), which, in the epidermis, is downregulated in activated keratinocytes (Waseem et al., 1999). We next investigated the expression of chondroitin sulphate proteoglycan 4 (CSPG4; also called melanoma chondroitin sulphate proteoglycan, MCSP). CSPG4 is known to facilitate cell proliferation and to protect against apoptosis (Chen et al., 2024). In human epidermis, CSPG4 expression defines a population of cycling stem cells (Ghali et al., 2004, Gunnarsson et al., 2017). Moreover, CSPG4 expression is upregulated in oral squamous cell carcinoma and strongly associated with loco-regional tumour relapse (Farnedi et al., 2015). The CSPG4 expression pattern we observed was opposite to that of NGFR. In NOM, CSPG4 expression was low but detectable at the top of the papillary lamina propria, similar to the situation in human interfollicular epidermis (Ghali et al., 2004, Gunnarsson et al., 2017). CSPG4 expression in RET-OLP was increased and even more so in ATR-OLP where all the basal cells and a few parabasal cells stained positive (Figure 3J–L, Table 1). Combined with our analysis of proliferation markers, the presence of NGFR/CSPG4 double-positive basal cells in the basal cell layer of RET-OLP epithelium (Figure 4A–C) is indicative of activation of those OESCs that remain undamaged by the immune cell attack. Consistent with its known role in promoting integrin activity (Yang et al., 2004), we observed co-localization of CSPG4 with ITGB1 in OLP tissue, including in patches of basal cells that had survived the immune cell attack (Figure 4D–F).

**Figure 4.**
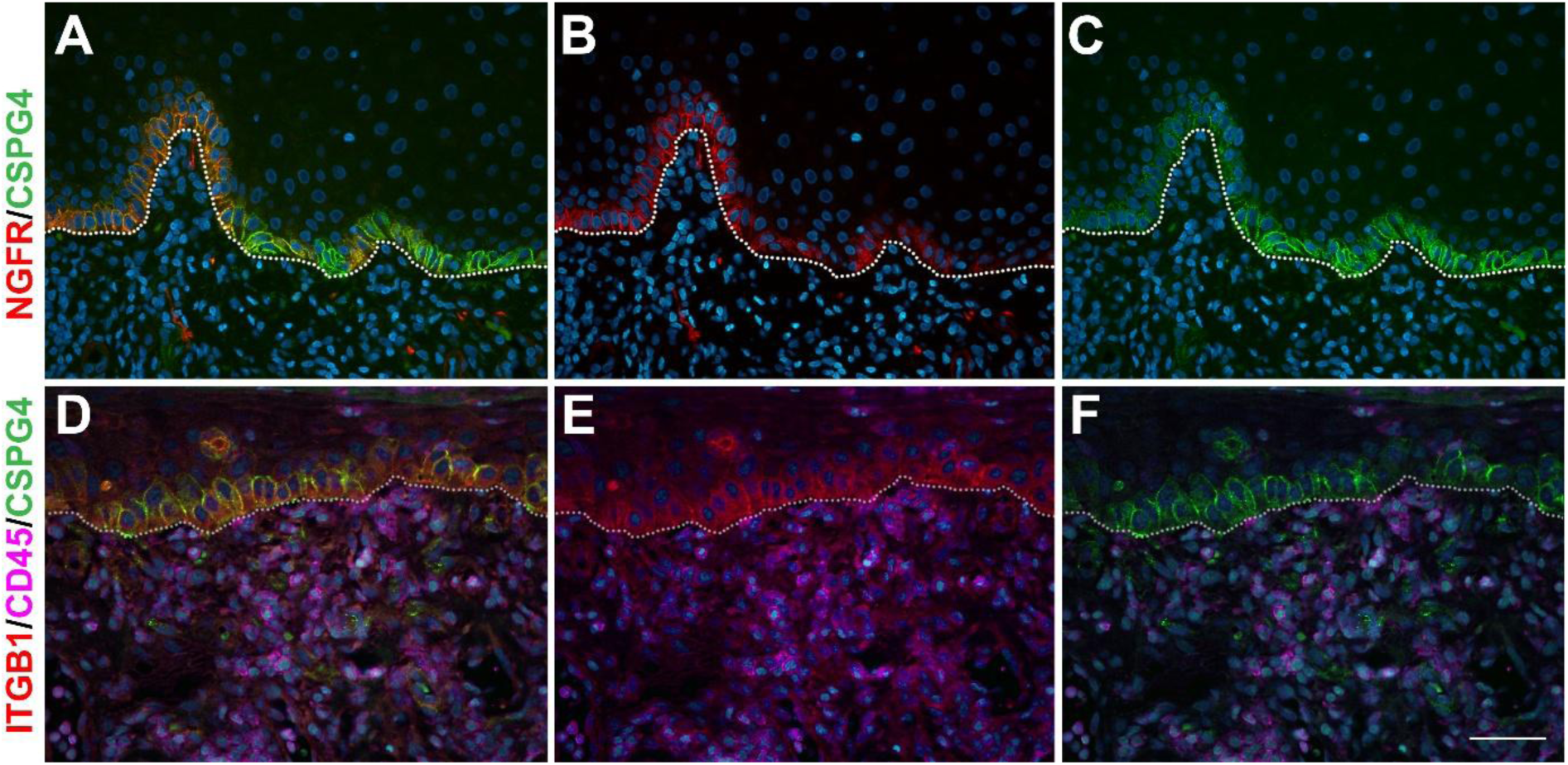
Activation of OESCs in OLP. Representative images show expression of NGFR and CSPG4 (A–C) in reticular OLP, and of ITGB1 and CSPG4 in atrophic OLP (D–F). Tissue sections were also labelled for DAPI (blue), and in (D–F) they were stained for CD45 (pink) to label immune cells. Dashed lines indicated the boundary between epithelium and underlying stromal tissue. Bar, 50 µm.

**Table 1.** Frequencies and intensities of staining for NGFR and CSPG4 in the basal cell layer of clinically normal oral mucosa (NOM), and reticular (RET-OLP) and atrophic oral lichen planus (ATR-OLP).

|  | NGFR |  |  | CSPG4 |  |  |
| --- | --- | --- | --- | --- | --- | --- |
|  | Moderate/<br>strong<br>staining of<br>all basal<br>cells | Restricted,<br>weak basal<br>cell staining | Negative<br>staining<br>of all<br>basal<br>cells | Positive staining<br>of basal cells at<br>the tips of the<br>papillary lamina<br>propria | Positive<br>staining of<br>some basal<br>cells | Positive<br>staining<br>of all<br>basal<br>cells |
| **NOM** | 12/12 | 00/12 | 00/12 | 06/12 | 05/12 | 01/12 |
| **RET-**<br>**OLP** | 03/11 | 06/11 | 02/11 | 00/11 | 03/11 | 08/11 |
| **ATR-**<br>**OLP** | 00/10 | 06/10 | 04/10 | 00/10 | 01/10 | 09/10 |

The transcriptional co-regulator protein YAP is characterised as a key regulator of epidermal stem cell self-renewal (De Rosa et al., 2019, Walko et al., 2017). In agreement with the findings from other groups (Hiemer et al., 2015, Omori et al., 2020), we found YAP to be strongest expressed in the basal cell compartment in NOM, where most cells also displayed nuclear YAP localisation (Figure 5A, Aii, Aiii). However, basal cells with nuclear YAP rarely stained positive for Ki-67 (Supplementary Figure S3), arguing against a role of YAP in driving OESC proliferation in NOM during homeostasis. In RET-OLP and ATR-OLP, some basal cells were able to maintain nuclear YAP localisation (Figure 5Biii, Ciii), but YAP expression was strongly reduced across the entire epithelium in ATR-OLP (Figure 5Cii). Altogether, our findings reveal marked epithelial tissue remodelling in OLP, which is linked to OESC activation, increased cell proliferation, and aberrant terminal differentiation.

**Figure 5.**
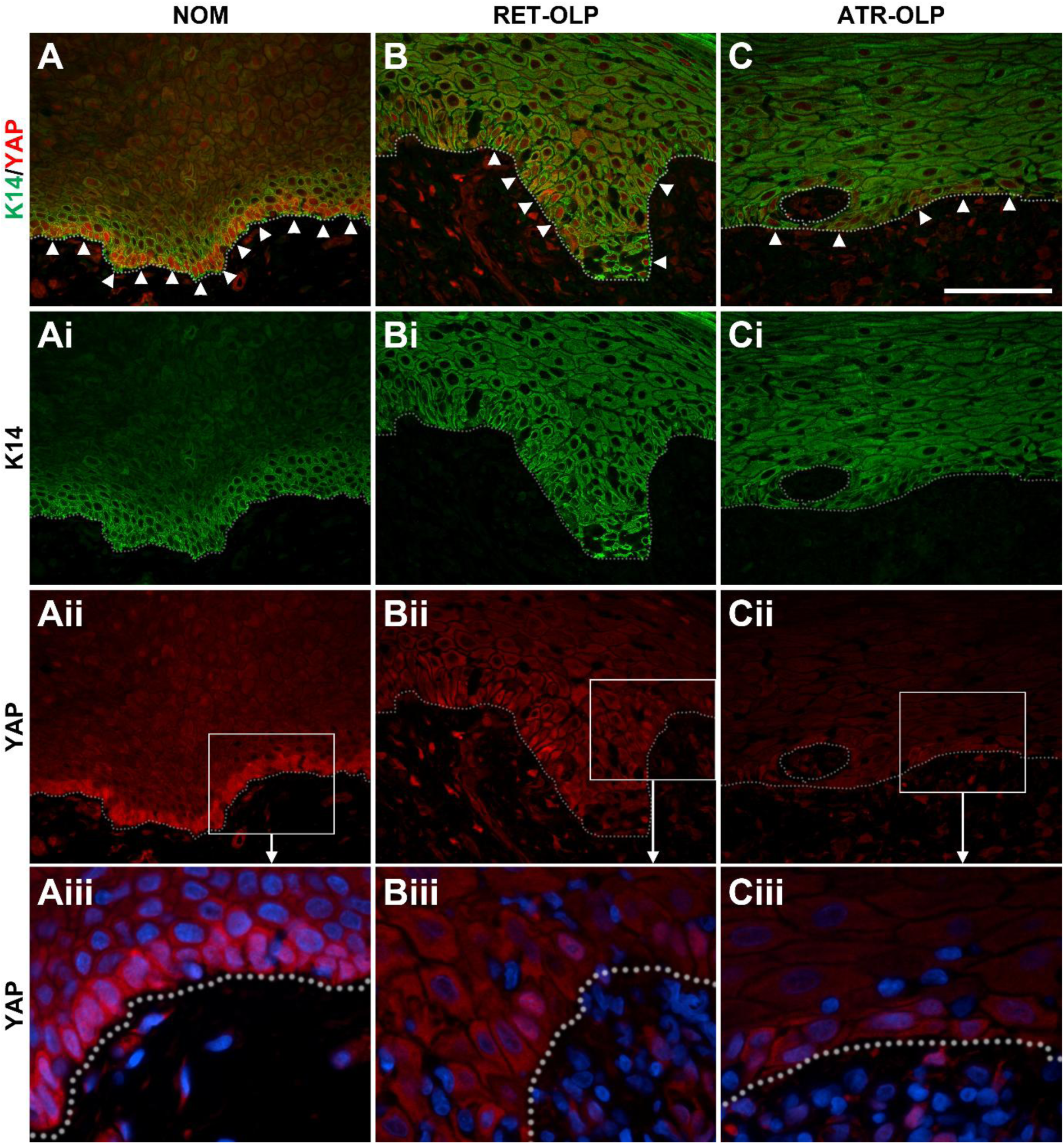
Altered expression of YAP in OLP. Representative images show expression of keratin (K)14 (green colour) (A–C, Ai–Ci) and of YAP (red colour) (A–C, Aii–Cii, Aiii–Ciii) in normal oral mucosa (NOM; A, Ai, Aii, Aiii), reticular OLP (RET-OLP; B, Bi, Bii, Biii), and atrophic OLP (ATR-OLP; C, Ci, Cii, Ciii). Dashed lines indicated the boundary between epithelium and underlying stromal tissue. In (Aiii–Ciii), cells were also labelled for DAPI (blue). Arrowheads in (A–C) indicate basal cells with nuclear YAP. Bar, 50 µm.

## Discussion

Here, we investigated the role that OESCs may play in OLP. In NOM, we identified OESCs in the basal epithelial cell layer as positive for the stem cell markers K15, K19, NGFR, ITGB1, and YAP, and negative for K4 and K16. In addition, OESCs are negative for p21. CSPG4, another stem cell marker (Gunnarsson et al., 2017), was also expressed, mostly at the top of the papillary lamina propria. The basal cell layer contained only ∼8% of all Ki-67-positive epithelial cells. The parabasal layer of NOM consists of 1-2 cell layers, has the profile of K5^+^/K8^+^/K14^+^/K15^+^/K4^-^/K16^-^/K19^-^/NGFR^-^, and parabasal cells comprise the majority of Ki-67-positive epithelial cells. In addition, many parabasal cells expressed p21. These findings are in line with studies of other types of human mucosal epithelia (esophagus, ectocervix, vagina) which showed that the fraction of proliferating cells is much lower in the basal cell layer compared to the parabasal epithelial cell layers (Andl et al., 2016, Dunaway et al., 2019, Harris et al., 2025). Incidentally, the numbers of proliferating cells presently observed in NOM were higher than observed in earlier reports. We used digital pathology for a survey of proliferation across a larger mucosal epithelial surface area, and this broader, non-selective approach, unlike prior work that selected restricted epithelial areas within mucosal biopsies, likely explains those higher numbers. Together, our findings are compatible with the existence of a one-layer basal cell compartment of ‘reserve’ OESCs in human oral mucosa. Most of these OESCs apparently cycle slowly as their majority is neither proliferating (Ki-67^-^), nor cell cycle arrested and differentiation-committed (p21^-^). The OESCs then give rise to a population of proliferating (Ki-67^+^) and ultimately terminal differentiation-committed (p21^+^) TACs in the parabasal cell layers. This task division between the slowly cycling OESCs and the frequently dividing and differentiating TACs is taken to be crucial for long-living species, since DNA replication in stratified squamous epithelial stem cells is thought to contribute to background mutation rate and thereby to cancer, taking into account the vast numbers of cell divisions that are taking place in these tissues (Dunaway et al., 2019).

The pathogenesis of OLP is still unclear. Without doubt, immune and/or inflammatory responses inflict damage to the epithelium, causing epithelial cell death, histologically seen as liquefactive degeneration in the basal cell layer. However, we observed that the stemness-associated protein ITGB1, a key component of focal adhesions, and the hemidesmosome component COL17A1, were largely preserved in significant areas of RET-OLP and ATR-OLP where the basal cell layer still occurred in a regular, organized fashion. This is consistent with previous observations that hemidesmosome homeostasis is disturbed in OLP but that the zones with intact or partially intact hemidesmosomes still can provide sufficient anchorage of the epithelium to the lamina propria to avoid blister formation (Schreurs et al., 2020a). Other stem cell- and differentiation characteristics in these zones were, however, significantly changed as compared with NOM. 1) The expression of the stem cell marker NGFR was markedly reduced (in RET-OLP) or absent (in ATR-OLP). Holoclone colonies (OESCs) established from the epithelium of NOM strongly express NGFR in their cell membrane, while meroclones (TACs with high proliferative capacity) show intermediate-, and paraclones (TACs with exhausted proliferative capacity) no expression of NGFR (Nakamura et al., 2007). This could mean that in OLP a vast majority of OESCs have left their quiescent state and are in an activated, proliferative state. The increased staining for CSPG4, a transmembrane proteoglycan found to be a useful marker for sorting actively cycling stem cells from human epidermis (Gunnarsson et al., 2017), also supports this notion. The presence of NGFR/CSPG4 double-positive basal cells in RET-OLP epithelium could mean that downregulation of NGFR and upregulation of CSPG4 represents a cell state switch between quiescence and activation. 2) Ki-67 expression was increased in the basal cell layer in OLP as compared with NOM. This shows that the OESCs in OLP are highly proliferating, probably in an attempt at restituting the epithelium that is exposed to the aggressive immune milieu. 3) Even though the OESCs in OLP start increased proliferation, normal mucosal homeostasis is not restored, as evident from the absence of the normal terminal differentiation programme, exemplified by reduced or absent suprabasal expression of KRT4. Instead, the progeny of the activated OESCs rapidly turn on expression of KRT16, and, at least in some epithelial areas, these cells commit to differentiation towards a keratinizing squamous epithelium. The presently observed changes of the terminal differentiation programme can explain the hyperkeratosis seen in OLP. 4) CSPG4 expression was increased in RET-OLP and even more so in ATR-OLP where all the basal cells and a few parabasal cells stained positive. This increased CSPG4 expression can be related to the secretion of IFNγ, TNFα, and IL1 in the microenvironment (Ampofo et al., 2017, Bonnet et al., 2011). CSPG4 can mediate protection from apoptosis induced by TNFα through activation of the PI3K/Akt signaling pathway (Chekenya et al., 2008, Yang et al., 2004). As the basal cells are under the pressure of a cell death-promoting milieu, CSPG4 upregulation can be a survival response of the basal cells against induction of cell death in OLP. Expression of p21 was also increased and this can also protect against apoptosis by binding and inhibiting pro-apoptotic proteins such as procaspase-3 and stress kinases (Wang et al., 2021). The observation of induction of protection against cell death aligns with other findings showing upregulation of anti-apoptotic factors, including FLIP_L_ and cIAP1/2 and increased expression of pro-apoptotic and anti-necroptotic FADD (Bonnet et al., 2011, Karatsaidis et al., 2007). Taken together, the OESCs in the zones without liquefaction degeneration appear to have left their state of quiescence, are actively dividing, are protecting themselves against untimely cell death, and proceed into a premature terminal differentiation program that might result in an atrophic epithelium.

Upregulated expression of CSPG4 may act as an oncoantigen, associated with aggressive SCC (Chen et al., 2022) and relapse of head and neck cancers (Farnedi et al., 2015). Also, in recessive dystrophic epidermolysis bullosa (RDEB), CSPG4 expression in skin precedes development of cutaneous SCC and promotes a more invasive tumour cell phenotype (Macaulay et al., 2024). The increased expression of CSPG4 in OLP might therefore contribute to the known increased risk for malignant transformation in the absence of the commonly recognised cancer-causing agents of the oro-pharynx (e.g. tobacco, alcohol and oncogenic types of human papilloma virus (Pimolbutr et al., 2024)).

In mouse epidermis, there is a clear positive correlation between the expression of proliferation markers and nuclear YAP localisation, and functional studies have confirmed YAP as a key regulator of epidermal keratinocyte proliferation during tissue development, homeostasis, and repair (Sedov et al., 2022, Walko et al., 2017, Zhang et al., 2011). However, we have shown here that in NOM, nuclear YAP localization, indicative of its activated state, is confined to the largely quiescent basal cell layer. This suggests that in NOM, and possibly also in other human squamous epithelia with compartmentalization into quiescent stem cell and proliferative TAC compartments, YAP functions to maintain the slowly cycling, undifferentiated population of stem cells with long-term regenerative capacity. The rapid replicative senescence of YAP-deficient human epidermal stells isolated from the skin of patients suffering from the blistering disease Junctional Epidermolysis Bullosa (JEB) supports this notion (De Rosa et al., 2019). The reduced YAP expression in JEB epidermis is linked to disturbed hemidesmosome assembly (De Rosa et al., 2019). Since hemidesmosome homeostasis is disturbed in ATR-OLP (Schreurs et al., 2020a)(Figure 3F), this may explain the presently observed reduced YAP expression. However, in RET-OLP and ATR-OLP some basal cells with nuclear YAP localisation were still detectable, highlighting that OESCs with long-term tissue regenerative capacity are not totally lost in OLP.

In conclusion, our findings highlight activation of slowly cycling OESCs that have retained their ability to enter the cell cycle upon the immune cell attack as a regenerative response driving tissue repair in OLP. The present results may underlie the clinical presentation and behaviour of oral lichen planus – by virtue of an intact means of cellular proliferation during immune attack, the oral epithelium is able to withstand lymphocytic attack. Hence disease predominantly manifests as epithelial hyperproliferation (e.g. reticular, papular or plaque-like OLP) and rarely as extensive areas of erosion (i.e. atrophy) or ulceration (Ingafou et al., 2006).

## Material and Methods

### Patient samples

Patients with OLP were recruited from the Oral Medicine unit of University College London Eastman Dental Hospital in London, UK, and from private dental practices in Oslo and Hamar, Norway. Written consent was obtained from all donors. Clinical diagnosis, gender, age, and smoking habits were recorded. The study was carried out with approval from the Regional Ethical Committees in the United Kingdom (Health Research Authority, Jarrow, Great Britain, 17/YH/0100) and Norway (REK Sørøst, Oslo, Norway, 2014/1724, 2021/347736 and 2022/290077) and was in accordance with the Declaration of Helsinki.

Histological evaluation was based on routinely H&E-stained formalin-fixed tissue specimens, according to established diagnostic criteria for oral lichen planus (Odell, 2025) and carried out by TMS. Specimens used in this study did not show epithelial dysplasia. Patients that were treated by systemic immunosuppressants, corticoids or biological modifiers, or with diagnosis of lichenoid reaction or GVHD, were excluded. Samples of normal, healthy, oral mucosa were obtained from clinically healthy buccal mucosa during third molar surgery or implant surgery. These samples were taken from the distal part of the releasing incision in the buccal mucosa.

All UK biopsies were fixed in 10 % neutral buffered formalin for 24–72 h, whereafter they were transferred to 70 % (v/v) ethanol and then stored at 4 °C. Prolonged cold storage in 70 % (v/v) ethanol did not affect staining intensity for any of the used markers as determined by test runs (not shown). Biopsies from Norway were placed in 10 % formalin and sent by regular mail to the Department of Pathology, The National Hospital, Oslo. All biopsies were dehydrated in graded ethanol, cleared in xylene and embedded in paraffin. The characteristics of the samples are summarized in Supplementary Table S2.

### Immunohistochemistry and immunofluorescence microscopy

Immunohistochemical processing and evaluation was done at the Institute of Oral Biology, University of Oslo. Four-micron thick paraffin sections were air dried and kept at 4 °C until further processing. Immediately before staining, sections were heated for 2 h at 60 °C, deparaffinized in two changes of xylene for 5 min each and rehydration through graded alcohols in 2 changes of absolute ethanol, 96% (v/v) and 70% (v/v) ethanol for 2 min each. Heat-induced antigen retrieval was performed by placing the slides in a Hellendahl jar filled with 0.05% (v/v) citraconic anhydride pH 7.4 (Sigma-Aldrich) using a decloaking chamber (Biocare medical) at 100 °C for 15 min and 90 °C for 10 seconds. The chamber was cooled down for 10 min on ice before the slides were rinsed with tap-water and equilibrated in phosphate-buffered saline (PBS). Sections were blocked for 1h with 5% (w/v) serum matching the source of the secondary antibody, prior to overnight incubation with primary antibodies at 4 °C and matching secondary antibodies for 2 h at room temperature (RT). Dilutions of sera and antibodies were made in antibody diluent containing 1% (w/v) bovine serum albumin in PBS with 0.05% ProClin 300 (v/v) (Sigma Aldrich) as preservative. Between incubation steps, sections were washed with PBS. Supplementary Table S3 presents a detailed list of the antibodies used. For multi-immunofluorescence staining, primary antibodies and secondary antibodies were mixed during their respective steps.

For immunofluorescence, the staining procedure continued with a 1h incubation at RT with fluorophore-labelled streptavidin containing 4’,6-diamidino-2-phenylindole (DAPI; Molecular Probes) before coverslips were mounted with polyvinyl alcohol mounting medium containing DABCO (Sigma-Aldrich). For chromogenic detections, sections were incubated with peroxidase-labeled Avidin-Biotin-Complex (Vectorlabs) for 30 min at RT, and signal was visualized using 3’ diaminobenzidine (DAB) as chromogen for 8 min and counterstained with Mayer’s hematoxylin (Dako).

As the staining pattern was different for primary antibodies of the same subclass, they served as control for non-specific binding of antibodies within similar subclasses. No unspecific binding was detected (Supplementary Figure S4).

Stained sections were imaged on a Nikon E90i epifluorescence microscope equipped with Ri1 and DS10 cameras using NIS-Elements Freeware v. and NIS-Elements BR-software v5.42 (Nikon). Evaluations were done using images taken with a 20x objective.

### Quantitative analysis of Ki-67 expression

A summary of the workflow is presented in Supplementary Figure S5. Sections stained for the proliferation marker Ki-67 were scanned using a 3DHistech panoramic MIDI slide scanner (3DHistech, Budapest, Hungary), followed by quantitative analysis using QuPath v0.5.1 (Bankhead et al., 2017). Scanned images were imported as brightfield (H-DAB setting) image type into a QuPath project. A line representing the basement membrane (BM) was drawn using the open polygon tool, just beneath the nuclei of the first aligned epithelial cell layer and the annotation classified as “BM” (Supplementary Figure S6, (1)). Basement membranes positioned within the epithelial area due to suboptimal tissue orientation were left unmarked. The epithelium was outlined with the closed polygon tool, covering the epithelial area excluding the superficial layer, and classified as “Epithelium” (Supplementary Figure S6, (2)). Zones with discontinuous or diffuse basement membrane were annotated separately and excluded (Supplementary Figure S6, (3)). Due to cutting level or suboptimal tissue orientation, discontinuous connective tissue papillae were sometimes observed within the epithelial annotation; these areas were annotated with the closed polygon tool and classified as “Ignore” (Supplementary Figure S6, (3)). The same procedure was used for indistinct/blurred areas or tissue folds. After selecting the areas to be analysed (Supplementary Figure S6, (4)), nuclei in the epithelial class were located using positive cell detection (*Analyze > Cell detection > Positive cell detections*) (Supplementary Figure S6, (5)). For image detection we used QuPath’s optical density sum method with a pixel size of 0.5 µm. Background radius for nuclear estimation was set to 8 µm and a Gaussian filter with Sigma = 2 µm was applied. The nuclear area was limited to 15-150 µm². The intensity threshold for nucleus detection was 0.1 with a maximum background intensity of 2; overlapping nuclei were split by shape. Detected nuclei were expanded by 5 µm to approximate the full cell area (including the nucleus), and boundaries were smoothed. To detect positive staining of the peroxidase substrate, the mean OD of nuclear DAB was used with a single threshold (default 0.2). Automatic detections were then corrected to match visual evaluation. First, nuclear detection was checked and, where necessary, parameters were slightly adjusted and detection rerun to correct the number of nuclei detected (Supplementary Figure S6, (6)). The hierarchy of the annotations was set (*Objects > Annotations > Resolve hierarchy*) to enable separate measurements for the exclusion areas. Second, the threshold for nuclear DAB detection was adjusted using an area containing both weakly and strongly stained nuclei while viewing DAB only (*View > Brightness/Contrast > DAB*) (Supplementary Figure S6, (7)). Then the detection classification was rerun (*Classify>Object classification>Set cell intensity classifications*), using the mean OD of nuclear DAB measurements. Thresholds for weak (1+), intermediate (2+) and strong (3+) positive cells were identified automatically. If the lowest threshold was manually adjusted, the intermediate was set to the midpoint of OD value between the weak and strong positive thresholds. Finally, the positions of detected nuclei within the epithelium were evaluated by analysing Distance to annotations (*Analyze > Spatial analysis > Distance to annotations 2D*) (Supplementary Figure S6, (8)). The detection measurement table was exported to Excel, and nuclei counted based on the intensity classification and distance to the nearest annotated basal membrane.

To obtain parameters enabling robust identification of the proliferative parabasal and basal cell layers, initial measurements were made on a set of randomly selected normal oral mucosa (NOM) biopsies stained for Ki-67. These revealed that the proliferative parabasal area in NOM comprised nuclei within 40 µm of the basement membrane, measured from the nucleus centre to the nearest BM annotation. Nuclei within 5 µm from the annotated BM were assigned to the basal cell layer. For measurement accuracy in both OLP and controls, the BM annotation was drawn directly beneath the nucleus, since first-layer cells in OLP may differ in shape from basal cells of healthy tissue.

We subsequently calculated the relative percentage of Ki-67-positive cells within the proliferative basal and parabasal areas.

### Bioinformatics analysis of published microarray data

Gene expression data from laser capture micro-dissected NOM and OLP epithelial tissues (Danielsson et al., 2014) were accessed through Gene Expression Omnibus (GSE52130). To identify genes that are differentially expressed between NOM and OLP tissues, we used the interactive web tool GEO2R (<u>GEO2R - GEO - NCBI</u>), which employs a variety of R packages from the <u>Bioconductor</u> project. Differential gene expression analysis was performed using *GEOquery* and *limma* (Davis and Meltzer, 2007), and applying limma precision weights (vooma). Multiple testing corrections were performed using the Benjamini & Hochberg false discovery rate method (p<0.05). Heatmaps were generated in R (v4.5.1).

### Statistical analysis and data presentation

Statistical analyses were performed with GraphPad Prism software (version 10.3.1) or R (v4.5.1). Statistical tests used to determine p-values are specified in Figure Legends.

## Funding information

The project was conducted with incentive funding from the University of Oslo and was additionally supported by the UK Research & Innovation (UKRI) Biotechnology and Biological Sciences Research Council (BBSRC, grant BB/T012978/1 to Gernot Walko), and by the National Institute for Health and Care Research (NIHR, UK) University College London Hospitals (UCLH) Biomedical Research Centre. Gernot Walko also acknowledges funding support from the Barts Charity (UK) Strategic Award to the Barts Centre for Squamous Cancer (Centre grant ‘A Centre of Excellence for Squamous Cancer‘, grant G-002030).

## Author contributions

<u>Olaf J. F. Schreurs</u>: Data Curation, Formal Analysis, Investigation, Methodology, Visualization, Writing: Original Draft Preparation, Writing: Review & Editing; <u>Stefano Fedele:</u> Conceptualization, Resources, Writing: Review & Editing; <u>Stephen Porter</u>: Conceptualization, Resources, Writing: Review & Editing; <u>Gry K. Kjølle</u>: Resources; <u>Karl Schenck</u>: Conceptualization, Formal Analysis, Funding Acquisition, Resources, Supervision, Writing: Original Draft Preparation, Writing: Review & Editing; <u>Tine M. Søland</u>: Conceptualization, Formal Analysis, Funding Acquisition, Resources, Supervision, Validation, Visualization, Writing: Review & Editing; <u>Gernot Walko</u>: Conceptualization, Formal Analysis, Funding acquisition, Project Administration, Resources, Supervision, Validation, Visualization, Writing: Original Draft Preparation, Writing: Review &Editing.

## Conflict of interest statement

The authors have no conflicts of interest to declare.

## Data availability

The authors declare that all data supporting the findings of this study are available within the paper and its supplementary information files. No novel sequencing data or code were generated for this study.

## Notes

### Competing Interest Statement

The authors have declared no competing interest.

## References

Ackerl R, Walko G, Fuchs P, Fischer I, Schmuth M, Wiche G. Conditional targeting of plectin in prenatal and adult mouse stratified epithelia causes keratinocyte fragility and lesional epidermal barrier defects. J Cell Sci 2007;120(Pt 14):2435–43.

Ampofo E, Schmitt BM, Menger MD, Laschke MW. The regulatory mechanisms of NG2/CSPG4 expression. Cell Mol Biol Lett 2017;22:4.

Andl CD, Le Bras GF, Loomans H, Kim AS, Zhou L, Zhang Y, et al. Association of TGFbeta signaling with the maintenance of a quiescent stem cell niche in human oral mucosa. Histochem Cell Biol 2016;146(5):539–55.

Andreasen JO. Oral lichen planus. 1. A clinical evaluation of 115 cases. Oral Surg Oral Med Oral Pathol 1968;25(1):31-42.

Bankhead P, Loughrey MB, Fernandez JA, Dombrowski Y, McArt DG, Dunne PD, et al. QuPath: Open source software for digital pathology image analysis. Sci Rep 2017;7(1):16878.

Bascones-Ilundain C, Gonzalez-Moles MA, Esparza G, Gil-Montoya JA, Bascones-Martinez A. Significance of liquefaction degeneration in oral lichen planus: a study of its relationship with apoptosis and cell cycle arrest markers. Clin Exp Dermatol 2007;32(5):556–63.

Bidarra M, Buchanan JA, Scully C, Moles DR, Porter SR. Oral lichen planus: a condition with more persistence and extra-oral involvement than suspected? J Oral Pathol Med 2008;37(10):582–6.

Boisnic S, Ouhayoun JP, Branchet MC, Frances C, Beranger JY, Le Charpentier Y, et al. Alteration of cytokeratin expression in oral lichen planus. Oral Surg Oral Med Oral Pathol Oral Radiol Endod 1995;79(2):207–15.

Bonnet MC, Preukschat D, Welz PS, van Loo G, Ermolaeva MA, Bloch W, et al. The adaptor protein FADD protects epidermal keratinocytes from necroptosis in vivo and prevents skin inflammation. Immunity 2011;35(4):572–82.

Byrd KM, Piehl NC, Patel JH, Huh WJ, Sequeira I, Lough KJ, et al. Heterogeneity within Stratified Epithelial Stem Cell Populations Maintains the Oral Mucosa in Response to Physiological Stress. Cell Stem Cell 2019;25(6):814–29 e6.

Carrozzo M, Porter S, Mercadante V, Fedele S. Oral lichen planus: A disease or a spectrum of tissue reactions? Types, causes, diagnostic algorhythms, prognosis, management strategies. Periodontol 2000 2019;80(1):105–25.

Chekenya M, Krakstad C, Svendsen A, Netland IA, Staalesen V, Tysnes BB, et al. The progenitor cell marker NG2/MPG promotes chemoresistance by activation of integrin-dependent PI3K/Akt signaling. Oncogene 2008;27(39):5182–94.

Chen K, Yong J, Zauner R, Wally V, Whitelock J, Sajinovic M, et al. Chondroitin Sulfate Proteoglycan 4 as a Marker for Aggressive Squamous Cell Carcinoma. Cancers (Basel) 2022;14(22).

Chen X, Habib S, Alexandru M, Chauhan J, Evan T, Troka JM, et al. Chondroitin Sulfate Proteoglycan 4 (CSPG4) as an Emerging Target for Immunotherapy to Treat Melanoma. Cancers (Basel) 2024;16(19).

Cheng YS, Gould A, Kurago Z, Fantasia J, Muller S. Diagnosis of oral lichen planus: a position paper of the American Academy of Oral and Maxillofacial Pathology. Oral Surg Oral Med Oral Pathol Oral Radiol 2016;122(3):332–54.

Cockburn K, Annusver K, Gonzalez DG, Ganesan S, May DP, Mesa KR, et al. Gradual differentiation uncoupled from cell cycle exit generates heterogeneity in the epidermal stem cell layer. Nat Cell Biol 2022;24(12):1692–700.

Conrad C, Gilliet M. Type I IFNs at the interface between cutaneous immunity and epidermal remodeling. J Invest Dermatol 2012;132(7):1759–62.

Danielsson K, Coates PJ, Ebrahimi M, Nylander E, Wahlin YB, Nylander K. Genes involved in epithelial differentiation and development are differentially expressed in oral and genital lichen planus epithelium compared to normal epithelium. Acta Derm Venereol 2014;94(5):526–30.

Davis S, Meltzer PS. GEOquery: a bridge between the Gene Expression Omnibus (GEO) and BioConductor. Bioinformatics 2007;23(14):1846–7.

de Pedro I, Galan-Vidal J, Freije A, de Diego E, Gandarillas A. p21CIP1 controls the squamous differentiation response to replication stress. Oncogene 2021;40(1):152–62.

De Rosa L, Secone Seconetti A, De Santis G, Pellacani G, Hirsch T, Rothoeft T, et al. Laminin 332-Dependent YAP Dysregulation Depletes Epidermal Stem Cells in Junctional Epidermolysis Bullosa. Cell Rep 2019;27(7):2036–49 e6.

Dunaway S, Rothaus A, Zhang Y, Luisa Kadekaro A, Andl T, Andl CD. Divide and conquer: two stem cell populations in squamous epithelia, reserves and the active duty forces. Int J Oral Sci 2019;11(3):26.

Eisen D. The evaluation of cutaneous, genital, scalp, nail, esophageal, and ocular involvement in patients with oral lichen planus. Oral Surg Oral Med Oral Pathol Oral Radiol Endod 1999;88(4):431–6.

Farnedi A, Rossi S, Bertani N, Gulli M, Silini EM, Mucignat MT, et al. Proteoglycan-based diversification of disease outcome in head and neck cancer patients identifies NG2/CSPG4 and syndecan-2 as unique relapse and overall survival predicting factors. BMC Cancer 2015;15:352.

Feldmeyer L, Suter VG, Oeschger C, Cazzaniga S, Bornstein MM, Simon D, et al. Oral lichen planus and oral lichenoid lesions - an analysis of clinical and histopathological features. J Eur Acad Dermatol Venereol 2020;34(2):e104–e7.

Fuchs E. Skin stem cells: rising to the surface. J Cell Biol 2008;180(2):273–84.

Ghali L, Wong ST, Tidman N, Quinn A, Philpott MP, Leigh IM. Epidermal and hair follicle progenitor cells express melanoma-associated chondroitin sulfate proteoglycan core protein. J Invest Dermatol 2004;122(2):433–42.

Gunnarsson AP, Christensen R, Praetorius J, Jensen UB. Isolating subpopulations of human epidermal basal cells based on polyclonal serum against trypsin-resistant CSPG4 epitopes. Exp Cell Res 2017;350(2):368–79.

Harris A, Burnham K, Pradhyumnan R, Jaishankar A, Hakkinen L, Gongora-Rosero RE, et al. Human-Specific Organization of Proliferation and Stemness in Squamous Epithelia: A Comparative Study to Elucidate Differences in Stem Cell Organization. Int J Mol Sci 2025;26(7).

Hedberg N, Ng A, Hunter N. A semi-quantitative assessment of the histopathology of oral lichen planus. J Oral Pathol 1986;15(5):268–72.

Hiemer SE, Zhang L, Kartha VK, Packer TS, Almershed M, Noonan V, et al. A YAP/TAZ-Regulated Molecular Signature Is Associated with Oral Squamous Cell Carcinoma. Mol Cancer Res 2015;13(6):957–68.

Iglesias-Bartolome R, Uchiyama A, Molinolo AA, Abusleme L, Brooks SR, Callejas-Valera JL, et al. Transcriptional signature primes human oral mucosa for rapid wound healing. Sci Transl Med 2018;10(451).

Ingafou M, Leao JC, Porter SR, Scully C. Oral lichen planus: a retrospective study of 690 British patients. Oral Dis 2006;12(5):463–8.

Janardhanam SB, Prakasam S, Swaminathan VT, Kodumudi KN, Zunt SL, Srinivasan M. Differential expression of TLR-2 and TLR-4 in the epithelial cells in oral lichen planus. Arch Oral Biol 2012;57(5):495–502.

Jones KB, Furukawa S, Marangoni P, Ma H, Pinkard H, D’Urso R, et al. Quantitative Clonal Analysis and Single-Cell Transcriptomics Reveal Division Kinetics, Hierarchy, and Fate of Oral Epithelial Progenitor Cells. Cell Stem Cell 2019;24(1):183–92 e8.

Jones KB, Klein OD. Oral epithelial stem cells in tissue maintenance and disease: the first steps in a long journey. Int J Oral Sci 2013;5(3):121–9.

Jones PH, Watt FM. Separation of human epidermal stem cells from transit amplifying cells on the basis of differences in integrin function and expression. Cell 1993;73(4):713–24.

Karatsaidis A, Hayashi K, Schreurs O, Helgeland K, Schenck K. Survival signalling in keratinocytes of erythematous oral lichen planus. J Oral Pathol Med 2007;36(4):215–22.

Karatsaidis A, Schreurs O, Axell T, Helgeland K, Schenck K. Inhibition of the transforming growth factor-beta/Smad signaling pathway in the epithelium of oral lichen. J Invest Dermatol 2003a;121(6):1283–90.

Karatsaidis A, Schreurs O, Helgeland K, Axell T, Schenck K. Erythematous and reticular forms of oral lichen planus and oral lichenoid reactions differ in pathological features related to disease activity. J Oral Pathol Med 2003b;32(5):275–81.

Kurzen N, Mubarak M, Eigemann J, Seiringer P, Wasserer S, Hillig C, et al. Death-Associated Protein Kinase 1 Dampens Keratinocyte Necroptosis and Expression of Inflammatory Genes in Lichen Planus. J Invest Dermatol 2025;145(8):1921–9 e13.

Kypriotou M, Huber M, Hohl D. The human epidermal differentiation complex: cornified envelope precursors, S100 proteins and the ‘fused genes’ family. Exp Dermatol 2012;21(9):643-9.

Li A, Kaur P. FACS enrichment of human keratinocyte stem cells. Methods Mol Biol 2005;289:87–96.

Li C, Tang X, Zheng X, Ge S, Wen H, Lin X, et al. Global Prevalence and Incidence Estimates of Oral Lichen Planus: A Systematic Review and Meta-analysis. JAMA Dermatol 2020;156(2):172–81.

Liu N, Matsumura H, Kato T, Ichinose S, Takada A, Namiki T, et al. Stem cell competition orchestrates skin homeostasis and ageing. Nature 2019;568(7752):344–50.

Macaulay ARK, Yang J, Price MA, Forster CL, Riddle MJ, Ebens CL, et al. Chondroitin sulfate proteoglycan 4 increases invasion of recessive dystrophic epidermolysis bullosa-associated cutaneous squamous cell carcinoma by modifying transforming growth factor-beta signalling. Br J Dermatol 2024;192(1):104–17.

Maciel GBM, Guse TL, Maciel RM, Danesi CC. Etiopathogenesis of Oral Lichen Planus: A Review. Head Neck Pathol 2025;19(1):73.

Maurizi E, Adamo D, Magrelli FM, Galaverni G, Attico E, Merra A, et al. Regenerative Medicine of Epithelia: Lessons From the Past and Future Goals. Front Bioeng Biotechnol 2021;9:652214.

Nakamura T, Endo K, Kinoshita S. Identification of human oral keratinocyte stem/progenitor cells by neurotrophin receptor p75 and the role of neurotrophin/p75 signaling. Stem Cells 2007;25(3):628–38.

Noske K, Stark HJ, Nevaril L, Berning M, Langbein L, Goyal A, et al. Mitotic Diversity in Homeostatic Human Interfollicular Epidermis. Int J Mol Sci 2016;17(2).

Odell EW. Cawson’s Essentials of Oral Pathology and Oral Medicine. 10 ed: Elsevier, 2025.

Omori H, Nishio M, Masuda M, Miyachi Y, Ueda F, Nakano T, et al. YAP1 is a potent driver of the onset and progression of oral squamous cell carcinoma. Sci Adv 2020;6(12):eaay3324.

Pan X, Hobbs RP, Coulombe PA. The expanding significance of keratin intermediate filaments in normal and diseased epithelia. Curr Opin Cell Biol 2013;25(1):47–56.

Pereira D, Sequeira I. A Scarless Healing Tale: Comparing Homeostasis and Wound Healing of Oral Mucosa With Skin and Oesophagus. Front Cell Dev Biol 2021;9:682143.

Pimolbutr K, Lim WT, Leeson R, Hopper C, Kalavrezos N, Liew C, et al. Prognosis of oral epithelial dysplasia in individuals with and without oral lichen planus. Oral Dis 2024;30(2):504–17.

Pontiggia L, Ahuja AK, Yosef HK, Rutsche D, Reichmann E, Moehrlen U, et al. Human Basal and Suprabasal Keratinocytes Are Both Able to Generate and Maintain Dermo-Epidermal Skin Substitutes in Long-Term In Vivo Experiments. Cells 2022;11(14).

Redmond CJ, Steiner SN, Cohen E, Johnson CN, Ozlu N, Coulombe PA. Keratin 15 promotes a progenitor cell state in basal keratinocytes of skin epidermis. J Cell Biol 2026;225(3).

Roopashree MR, Gondhalekar RV, Shashikanth MC, George J, Thippeswamy SH, Shukla A. Pathogenesis of oral lichen planus--a review. J Oral Pathol Med 2010;39(10):729–34.

Schlosser BJ. Lichen planus and lichenoid reactions of the oral mucosa. Dermatol Ther 2010;23(3):251–67.

Schluter H, Paquet-Fifield S, Gangatirkar P, Li J, Kaur P. Functional characterization of quiescent keratinocyte stem cells and their progeny reveals a hierarchical organization in human skin epidermis. Stem Cells 2011;29(8):1256–68.

Schreurs O, Balta MG, Karatsaidis A, Schenck K. Composition of hemidesmosomes in basal keratinocytes of normal buccal mucosa and oral lichen planus. Eur J Oral Sci 2020a;128(5):369–78.

Schreurs O, Karatsaidis A, Balta MG, Grung B, Hals EKB, Schenck K. Expression of keratins 8, 18, and 19 in epithelia of atrophic oral lichen planus. Eur J Oral Sci 2020b;128(1):7-17.

Sedov E, Koren E, Chopra S, Ankawa R, Yosefzon Y, Yusupova M, et al. THY1-mediated mechanisms converge to drive YAP activation in skin homeostasis and repair. Nat Cell Biol 2022;24(7):1049–63.

Sehbai A, Hamid MA, Ibrahim Z. Pembrolizumab-Induced Lichen Planus in a Patient With Non-Small-Cell Lung Carcinoma (NSCLC) That Correlates to Therapeutic Response. Cureus 2022;14(11):e31454.

Silverman S, Jr., Gorsky M, Lozada-Nur F. A prospective follow-up study of 570 patients with oral lichen planus: persistence, remission, and malignant association. Oral Surg Oral Med Oral Pathol 1985;60(1):30–4.

Thorn JJ, Holmstrup P, Rindum J, Pindborg JJ. Course of various clinical forms of oral lichen planus. A prospective follow-up study of 611 patients. J Oral Pathol 1988;17(5):213-8.

Thornhill MH. Immune mechanisms in oral lichen planus. Acta Odontol Scand 2001;59(3):174–7.

Topley GI, Okuyama R, Gonzales JG, Conti C, Dotto GP. p21(WAF1/Cip1) functions as a suppressor of malignant skin tumor formation and a determinant of keratinocyte stem-cell potential. Proc Natl Acad Sci U S A 1999;96(16):9089–94.

Walko G, Woodhouse S, Pisco AO, Rognoni E, Liakath-Ali K, Lichtenberger BM, et al. A genome-wide screen identifies YAP/WBP2 interplay conferring growth advantage on human epidermal stem cells. Nat Commun 2017;8:14744.

Wang L, Han H, Dong L, Wang Z, Qin Y. Function of p21 and its therapeutic effects in esophageal cancer. Oncol Lett 2021;21(2):136.

Wang S, Drummond ML, Guerrero-Juarez CF, Tarapore E, MacLean AL, Stabell AR, et al. Single cell transcriptomics of human epidermis identifies basal stem cell transition states. Nat Commun 2020;11(1):4239.

Waseem A, Dogan B, Tidman N, Alam Y, Purkis P, Jackson S, et al. Keratin 15 expression in stratified epithelia: downregulation in activated keratinocytes. J Invest Dermatol 1999;112(3):362–9.

Wiedemann J, Billi AC, Bocci F, Kashgari G, Xing E, Tsoi LC, et al. Differential cell composition and split epidermal differentiation in human palm, sole, and hip skin. Cell Rep 2023;42(1):111994.

Yang J, Price MA, Neudauer CL, Wilson C, Ferrone S, Xia H, et al. Melanoma chondroitin sulfate proteoglycan enhances FAK and ERK activation by distinct mechanisms. J Cell Biol 2004;165(6):881–91.

Yang Z, Deng M, Ren L, Fan Z, Yang S, Liu S, et al. Pyroptosis of oral keratinocyte contributes to energy metabolic reprogramming of T cells in oral lichen planus via OPA1-mediated mitochondrial fusion. Cell Death Discov 2024;10(1):408.

Zhang H, Pasolli HA, Fuchs E. Yes-associated protein (YAP) transcriptional coactivator functions in balancing growth and differentiation in skin. Proc Natl Acad Sci U S A 2011;108(6):2270–5.

Zhang Y, Bailey D, Yang P, Kim E, Que J. The development and stem cells of the esophagus. Development 2021;148(6).

