## Supplementary Figures and Tables for "The stem cell compartment in human oral mucosa and its activation in oral lichen planus"

**Supplementary data**

### **Supplementary Figure 1. Expression of p21 and Ki-67 in NOM and OLP mucosal epithelium.** Representative images show expression of Ki-67 (green colour) (A–C, Ai–Ci) and of p21 (red colour) (A–C, Aii–Cii) in normal oral mucosa (NOM; A, Ai, Aii), reticular OLP (RET-OLP; B, Bi, Bii), and atrophic OLP (ATR-OLP; C, Ci, Cii). Dashed lines indicated the boundary between epithelium and underlying stromal tissue. Arrowheads in (Aii–Cii) indicate p21-positive cells in the basal cell layer. Bar, 50 µm.

**Supplementary Figure 2. Expression levels of cytokeratin- and terminal differentiation marker mRNAs in OLP versus NOM epithelium.** (A) Significantly enriched or depleted RNAs in laser capture micro-dissected OLP epithelial tissue identified by differential gene expression analysis. Volcano plot shows statistical significance (-log 10 (adj. p-value)) versus relative RNA abundance (log2-fold change) in OLP compared to NOM epithelium. Red and blue colour: RNAs that passed the cut-offs for differential expression (log2-fold change>±1) and statistical significance (p<0.05). (B, C) Heatmaps show the log2-fold change in expression of selected keratin genes (B) and genes belonging to the epidermal differentiation complex (EDC)(C) in OLP versus NOM epithelium.

### **Supplementary Figure 3. Nuclear YAP localization does not indicate proliferating mucosal epithelial cells.** Expression of YAP (red colour) and Ki-67 (green colour) in normal oral mucosa (NOM; A), reticular OLP (RET-OLP; B), and atrophic OLP (ATR-OLP; C). Arrowheads in (A, B) indicate basal cells with nuclear YAP. Bar, 50 µm.

### **Supplementary Figure 4. Antibody controls for immunohistochemistry.** Negative control staining was performed on buccal mucosa of healthy individuals (NOM), and patients with reticular (RET-OLP) and atrophic (ATR-OLP) oral lichen planus by using control mouse IgG1, control mouse IgG2b, or pre-immune rabbit immunoglobulin, matching the highest concentration used for the respective antibodies.

### **Supplementary Figure 5. Pipeline for semi-automated analysis of Ki-67 expression.**

### **Supplementary Figure 6. Example of semi-automated analysis of Ki-67 expression.** Sequential analysis steps are numbered. For details, please refer to Materials and Methods section.

### **Supplementary Table 1. Summary of cytokeratin expression pattern in NOM versus OLP oral mucosa.**

### **Supplementary Table 2. Demographics of the volunteers of the study.**

### **Supplementary Table 3. Antibodies used in this study.**

**Supplementary Figure S1**

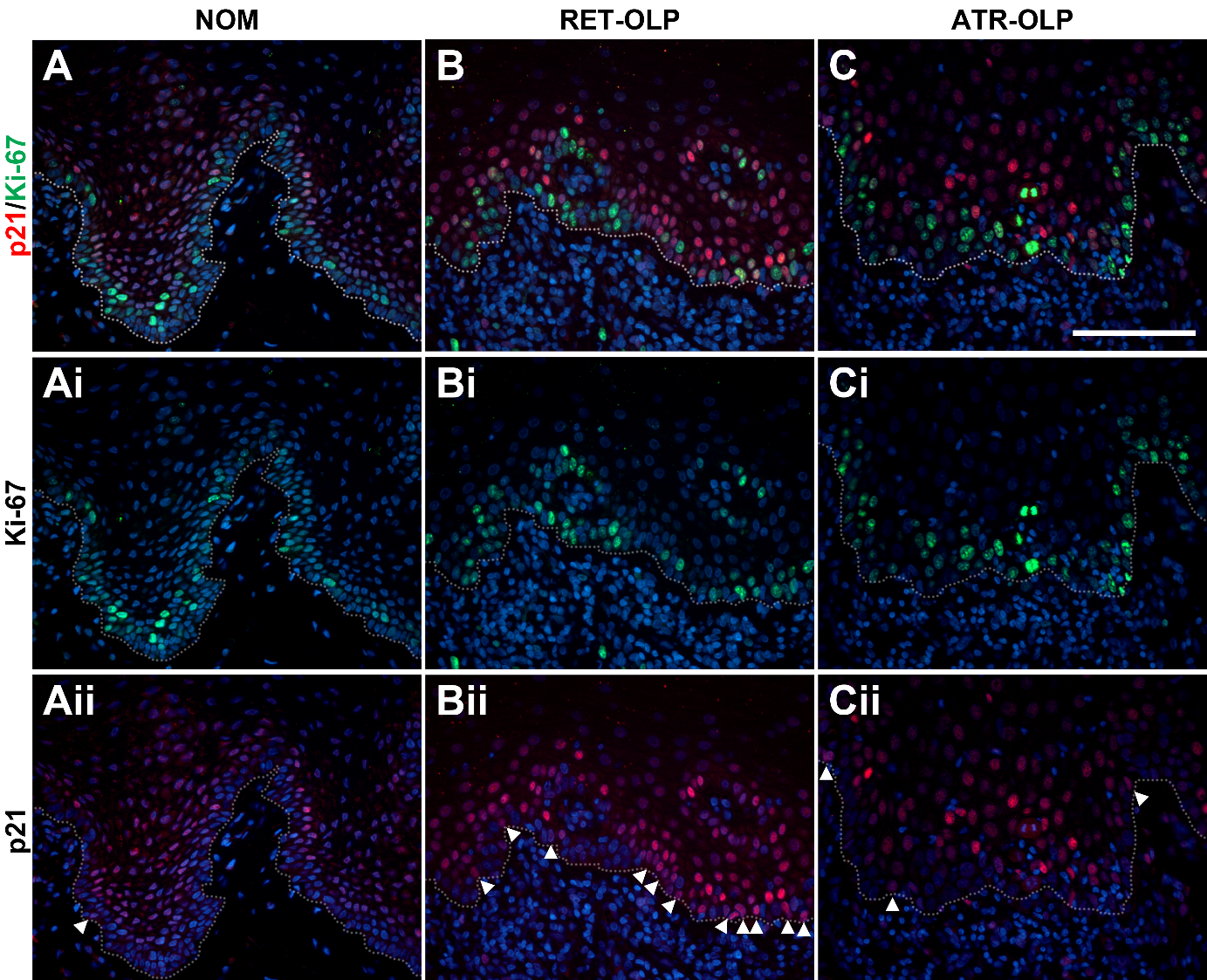

**Supplementary Figure S2**

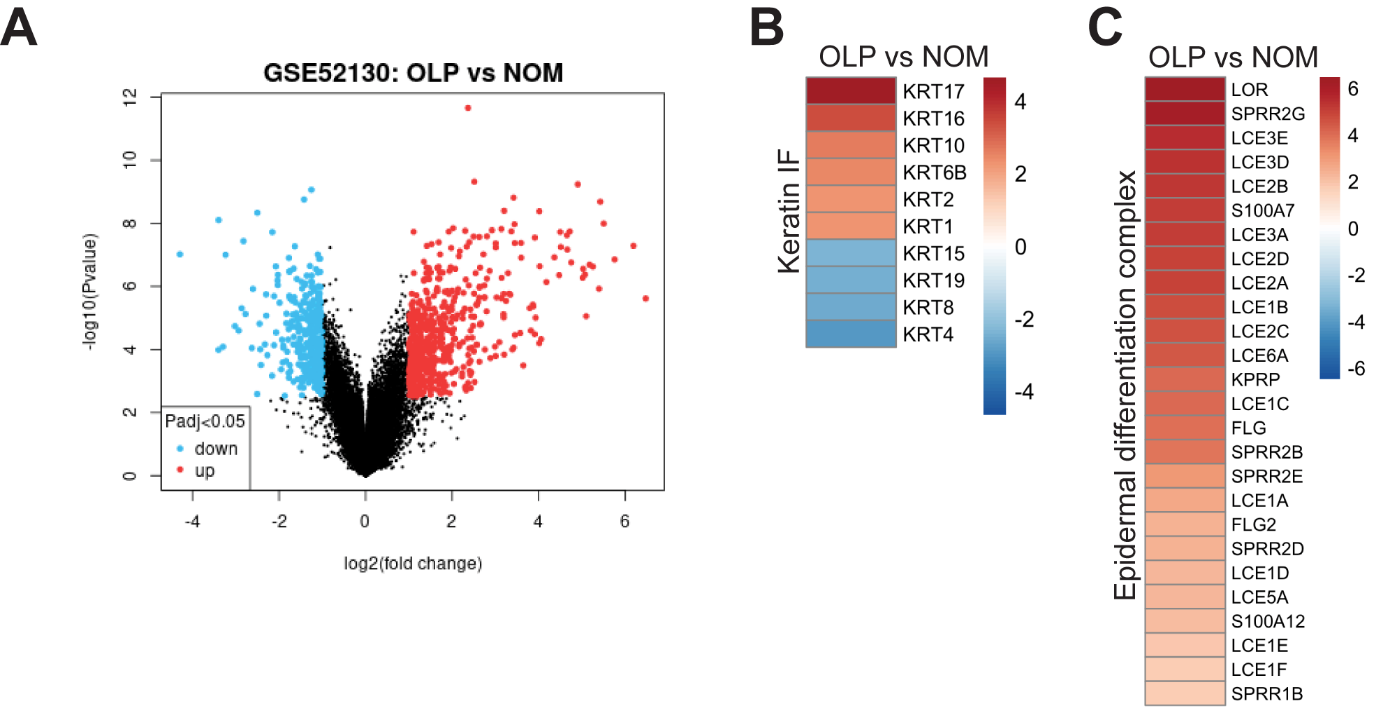

**Supplementary Figure S3**

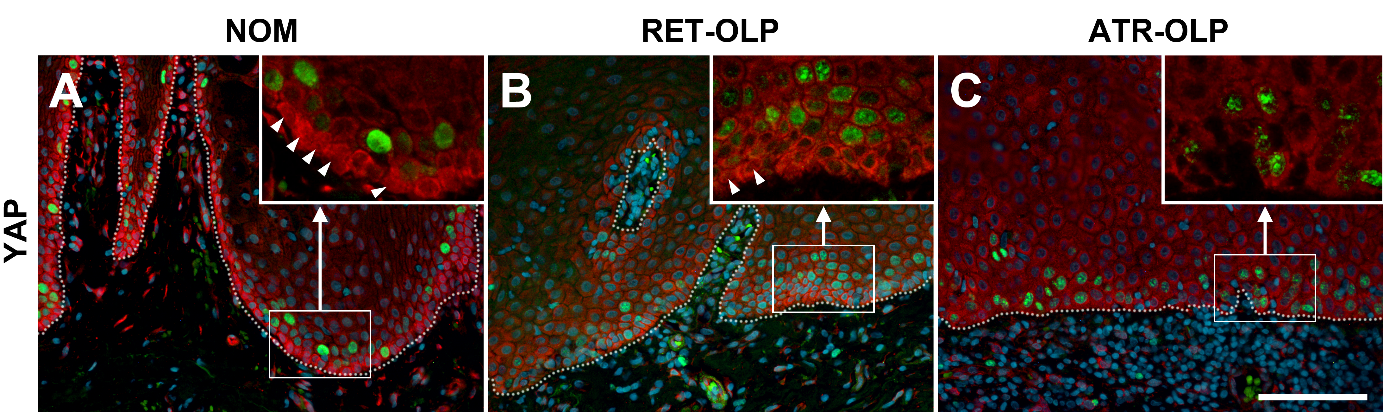

**Supplementary Figure S4**

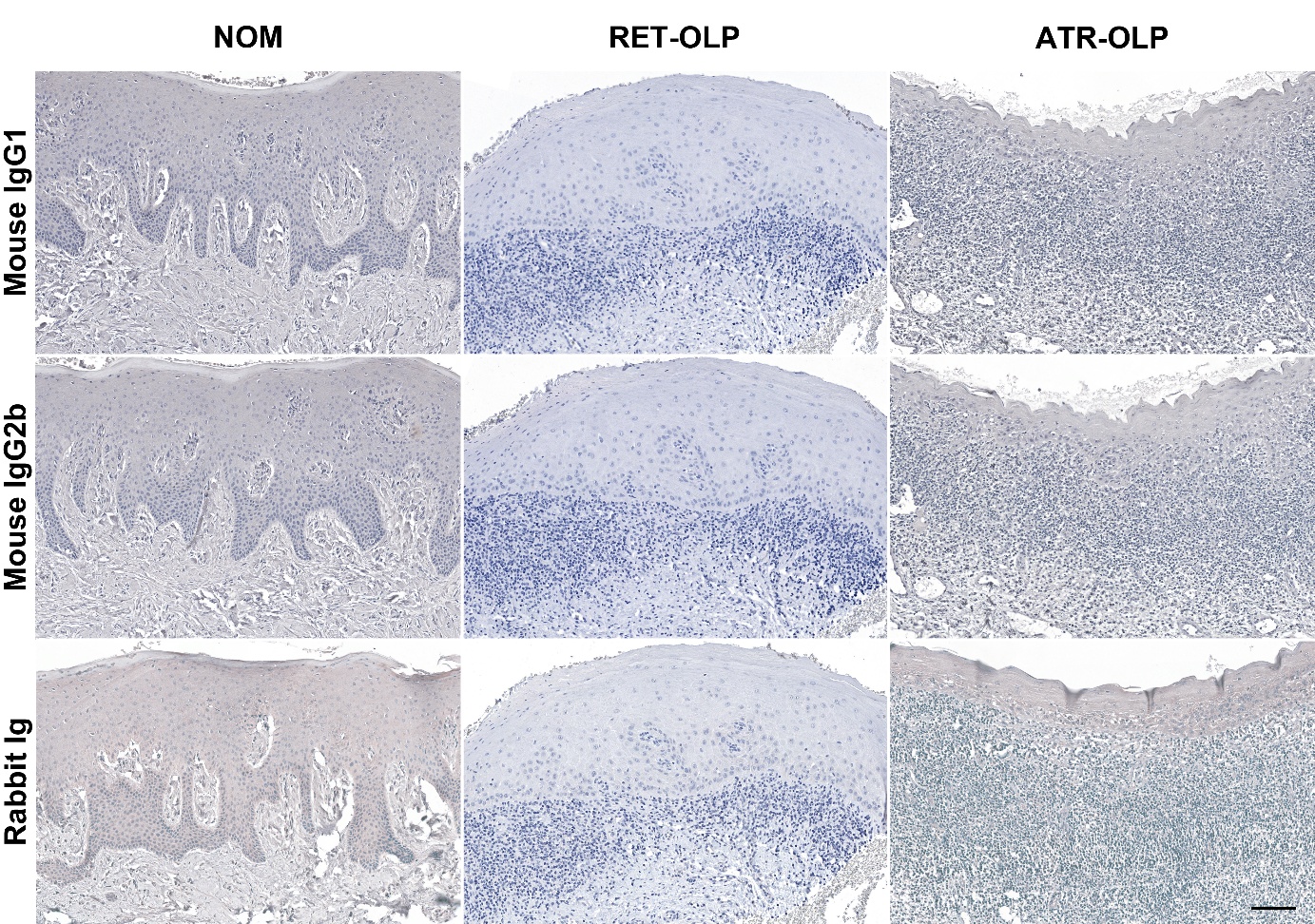

**Supplementary Figure S5**

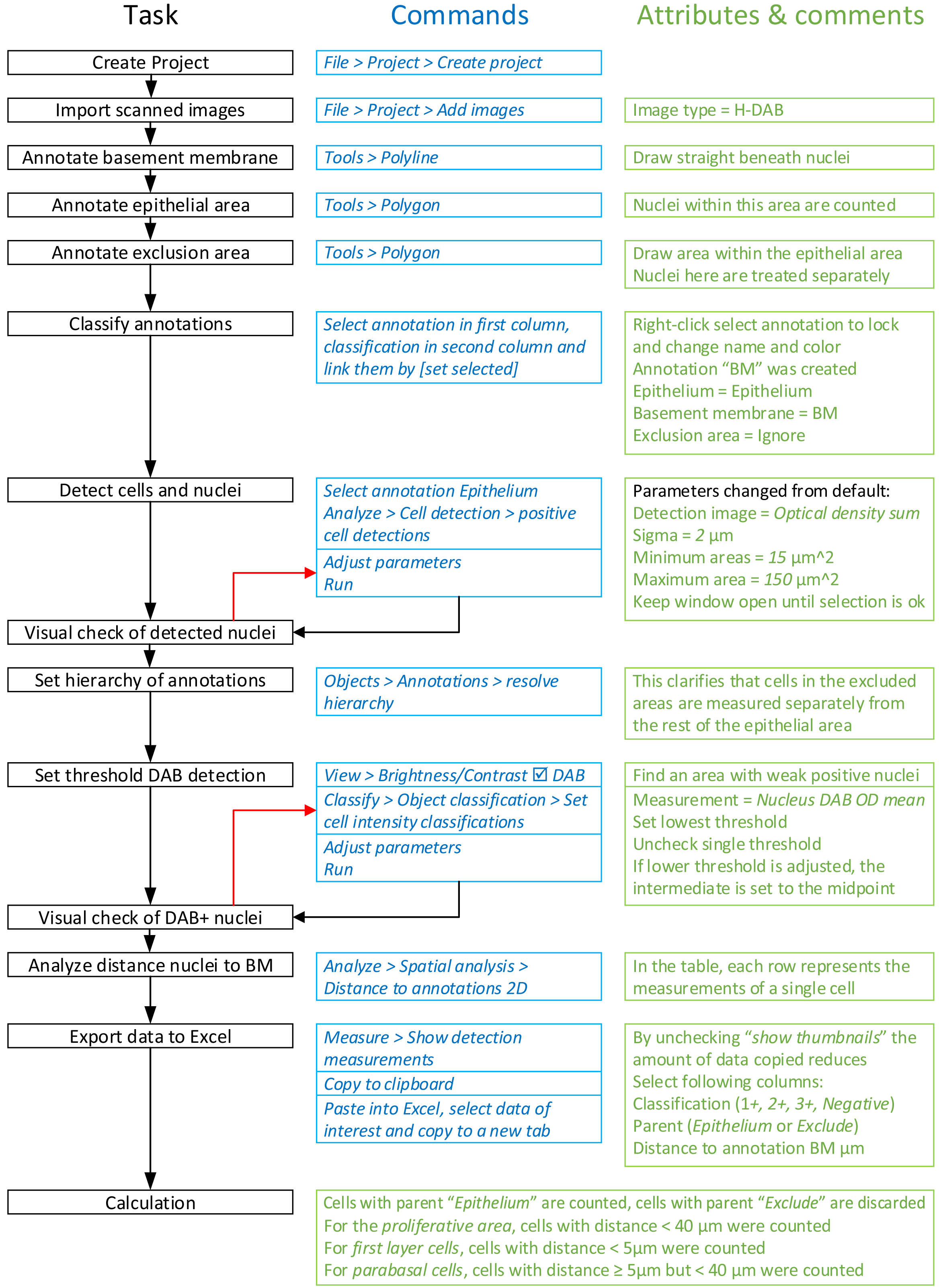

**Supplementary Figure S6**

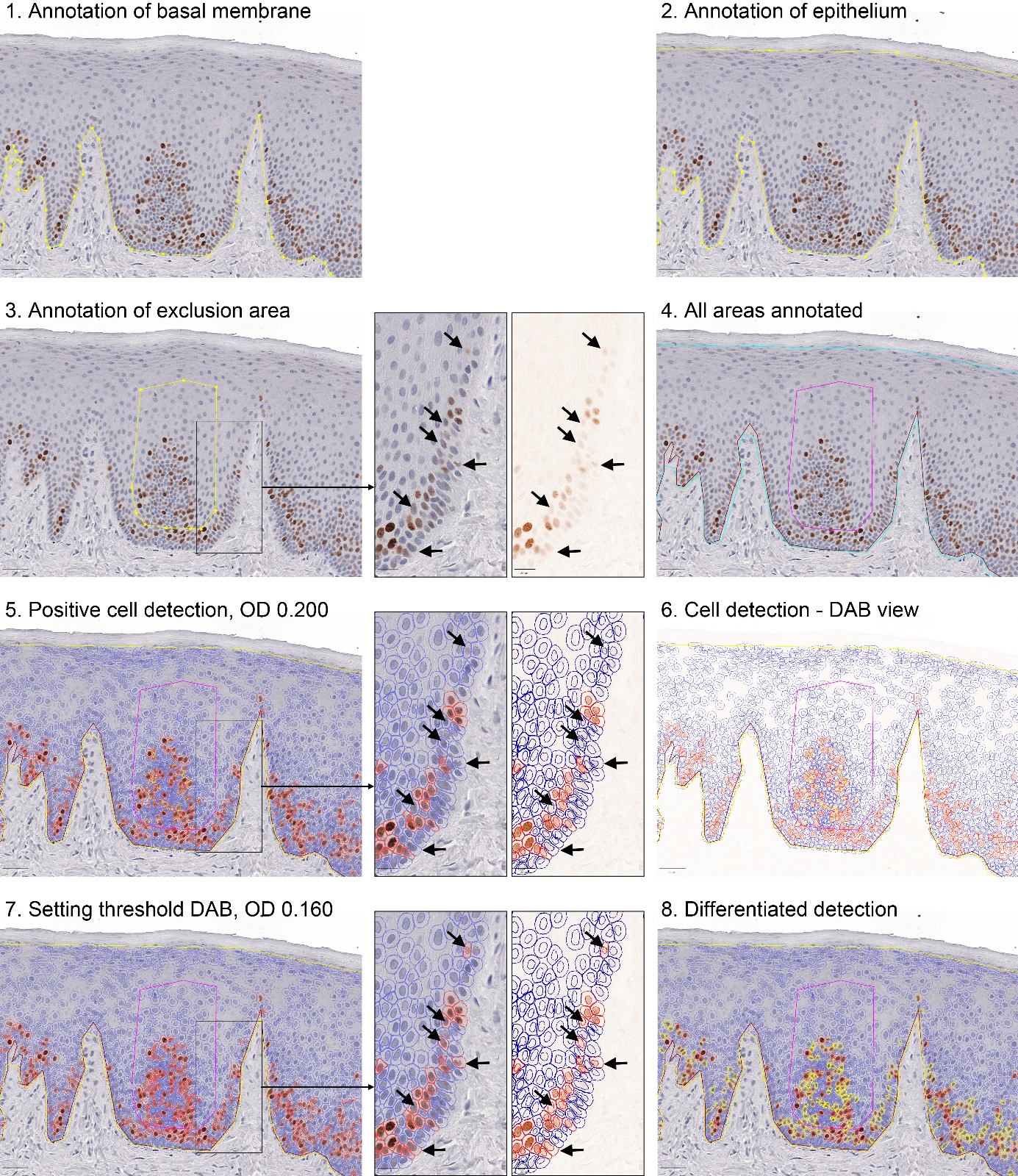

**Supplementary Table S1**

*Cytokeratin (K) staining in normal oral mucosa (NOM),*

*and reticular (RET-OLP) and atrophic (ATR-OLP) oral lichen planus.*

| Keratin | NOM | RET-OLP | ATR-OLP |
| --- | --- | --- | --- |
| K4 | Suprabasal terminally differentiating cells | Reduced or absent | As in RET-OLP |
| K5 | All cell layers, parabasal layer strongest | Expanded to the suprabasal cell layers | As in RET-OLP |
| K8 | Basal and parabasal layers | Patches of basal cells | Few basal cells |
| K10 | Absent or weak in suprabasal cells | Variable among cases, moderate to strong in suprabasal cells | As in RET-OLP, but with areas of continuous staining |
| K14 | Basal and first parabasal layer | Expanded to the suprabasal cell layers | As in RET-OLP |
| K15 | Basal and first parabasal layer | Patches of basal cells | Few basal cells |
| K16 | Non-proliferating cells above the parabasal cell layer | Some basal cells, most parabasal cells, cells above the parabasal cell layer | As in RET-OLP |
| K19 | Basal and some parabasal cells | No staining | As in RET-OLP |

**Supplementary Table S2**

*Demographics of the volunteers of the study*

| Variable | Persons with normal oral mucosa  (NOM) | Patients with reticular oral lichen planus  (RET-OLP) | Patients with atrophic oral lichen planus  (ATR-OLP) |
| --- | --- | --- | --- |
| *n* | 12 | 11 | 10 |
| Age (yr, range) | 40 (21 – 71) | 51 (34 – 71) | 61,5 (35 – 83) |
| Gender (male/female) | 8/4 | 2/9 | 2/8 |
| Smoker/non-smoker | 0/12 | 0/11 | 2/10 |

Values are given as *n*, *n/n*, or median (range)

**Supplementary Table S3**

*Antibodies used in this study*

| Antigen | Species | Vendor |
| --- | --- | --- |
| CD29, ITGB1 | Ms mAb | Lab Vision Cat# MS-596-P, RRID:AB_142776 |
| CD45 | Ms mAb | Diatec Cat# CD45 clone EO-1 |
| COL17A1 | Rb pAb | Sigma-Aldrich Cat# HPA043673, RRID:AB_10960893 |
| CSPG4 | Rb mAb | Cell Signaling Technology Cat# 43916, RRID:AB_3086773 |
| Cytokeratin 4 | Ms mAb | Santa Cruz Biotechnology Cat# sc-52321, RRID:AB_2249751 |
| Cytokeratin 5 | Rb mAb | DSHB Cat# CPTC-KRT5-2, RRID:AB_2820263 |
| Cytokeratin 8 | Rb mAb | Thermo Fisher Scientific Cat# MA5-14476, RRID:AB_10985243 |
| Cytokeratin 10 | Ms mAb | Santa Cruz Biotechnology Cat# sc-53252, RRID:AB_629835 |
| Cytokeratin 14 | Ms mAb | Bio-Rad Cat# MCA890, RRID:AB_322015 |
| Cytokeratin 15 | Ms mAb | Thermo Fisher Scientific Cat# MA5-11344, RRID:AB_10999819 |
| Cytokeratin 16 | Ms mAb | Thermo Fisher Scientific Cat# MA5-13730, RRID:AB_10983103 |
| Cytokeratin 19 | Ms mAb | Thermo Fisher Scientific Cat# MA5-12663, RRID:AB_10984317 |
| Ki-67 | Rb mAb | Thermo Fisher Scientific Cat# MA5-14520, RRID:AB_10979488 |
| Ki-67 | Ms mAb | Agilent Cat# M7240, RRID:AB_2142367 |
| NGFR | Ms mAb | Santa Cruz Biotechnology Cat# sc-13577, RRID:AB_627879 |
| p21 | Rb mAb | Abcam Cat# ab109520, RRID:AB_10860537 |
| YAP | Rb mAb | Cell Signaling Technology Cat# 14074, RRID:AB_2650491 |
| Mouse IgG1 control | Ms mAb | Agilent Cat# X0931, RRID:AB_2889134 |
| Mouse IgG2b control | Ms mAb | Agilent Cat# X0944 |
| Mouse IgG-Bio | Hrs pAb | Vector Laboratories Cat# BA-2000, RRID:AB_2313581 |
| Rabbit Ig control | Rb pAb | Eurogentec pre-immune serum ZNO06009/rb3166 |
| Rabbit IgG-AF488+ | Dk pAb | Thermo Fisher Scientific Cat# A32790, RRID:AB_2762833 |
| Rabbit IgG-Bio | Gt pAb | Vector Laboratories Cat# BA-1000, RRID:AB_2313606 |
| Rabbit IgG-Cy3 | Dk pAb | Jackson ImmunoResearch Labs Cat# 711-165-152, RRID:AB_2307443 |
| Mouse IgG1-AF488 | Gt pAb | SouthernBiotech Cat# 1070-30, RRID:AB_2794420 |
| Mouse IgG1-AF647 | Gt pAb | Thermo Fisher Scientific Cat# A-21240, RRID:AB_2535809 |
| Mouse IgG1-Bio | Gt pAb | Caltag Cat# M32115 |
| Mouse IgG1-Cy3 | Gt pAb | Jackson ImmunoResearch Labs Cat# 115-165-205, RRID:AB_2338694 |
| Mouse IgG2a-AF555 | Gt pAb | SouthernBiotech Cat# 1080-32, RRID:AB_2794491 |
| Mouse IgG2b-AF555 | Gt pAb | SouthernBiotech Cat# 1090-32, RRID:AB_2794533 |
| Mouse IgG2b-AF488 | Gt pAb | Thermo Fisher Scientific Cat# 2432059, RRID:AB_2535778 |
| Mouse IgG3-AF488 | Gt pAb | Jackson ImmunoResearch Labs Cat# 115-545-209, RRID:AB_2338858 |
